# Lateral flow devices can fast-track rabies virus genomic surveillance

**DOI:** 10.64898/2026.10.12.758520

**Authors:** Bea Crisostomo, Criselda Bautista, Mary Elizabeth Miranda, Daria L. Manalo, Shynie Vee. M. Telmo, Gian Rei Mangcucang, Sheryl Pablo, Michael Ralph Tañedo, Eleanor M. Rees, Sandeep Kasaragod, Nai Rui Chng, Katie Hampson, Kirstyn Brunker

## Abstract

Viral genomic data can resolve rabies transmission and guide outbreak response, but surveillance is often limited less by sequencing capacity than by getting suitable specimens from remote collection sites to a laboratory, a process that can take months. Here, we show that lateral flow devices (LFDs) used for rabies diagnosis can also serve as genomic specimens, integrating diagnosis and genomic sampling into a single workflow without additional specimen collection. We evaluated LFDs and FTA cards for rabies virus genome recovery under laboratory and field conditions. During surveillance across the island provinces of MIMAROPA, Philippines, the Bionote LFD had 99.1% sensitivity and 100% specificity against the gold-standard direct fluorescent antibody test (n = 319), with results available a median of 1 day after sampling versus 5 days for DFAT confirmation. Under laboratory conditions simulating field storage (ambient temperature or 37°C for up to 21 days), both residual LFD material and FTA cards supported near-complete genome sequencing, although genome recovery declined with storage duration, particularly at elevated temperatures. Consistent with this, field-collected LFDs yielded near-complete genomes up to 23 days of ambient storage but only partial genomes after 29-77 days, indicating a practical window of around three weeks for use. Repurposing LFDs already embedded in diagnostic workflows offers a practical route to decentralised genomic surveillance without additional sample collection or a cold chain. FTA cards provide a complementary preservation option, while further optimisation of specimen preparation, extraction and storage could improve genome recovery and extend the sequencing window.

## Introduction

Rabies is an invariably fatal but vaccine-preventable disease caused by the rabies virus (RABV), a single-stranded RNA virus, transmitted primarily through the saliva of infected animals, usually following bites or scratches (WHO 2018a). Domestic dogs account for more than 99% of human infections with rabies tens of thousands of human deaths each year. These are predominantly in low- and middle-income countries (Hampson et al. 2015, 2019). Mass dog vaccination and post-exposure prophylaxis can prevent these deaths, making the global target of eliminating dog-mediated rabies by 2030 achievable in principle, but reaching it will require that these interventions are guided by effective surveillance approaches (WHO 2018b).

Genomic surveillance can complement conventional case-based rabies surveillance by resolving transmission pathways, tracking viral movement and characterising circulating lineages (Yuson et al. 2024; Lushasi et al. 2022; Bourhy et al. 2016; Dellicour et al. 2019; Brunker et al. 2015, 2018; Gibson et al. 2022; Brunker et al. 2026). For example, phylogenomic analysis of the rabies epidemic in Arequipa, Peru, traced the dominant lineage to an introduction approximately three years before transmission was first detected (Brunker et al. 2026), and in the Philippines, integrating genomic and epidemiological data identified the likely human-mediated introduction of RABV to the previously rabies-free island of Tablas from mainland Luzon (Yuson et al. 2024). Realising this value for outbreak response, however, depends on sequence data being generated quickly enough to act on.

In many rabies-endemic settings, diagnosis and sequencing are centralised. Samples from suspected rabid animals are transported from peripheral collection sites to regional or national laboratories for confirmatory testing, usually with the direct fluorescent antibody test (DFAT). Subsequently, selected positive samples may then be referred for sequencing. This multi-step pathway can be slow, particularly where populations are geographically dispersed and transport infrastructure is limited. In the Philippines, for example, samples tested with lateral flow devices (LFDs) at the point of collection in MIMAROPA, a region of dispersed island provinces where rabies has recently re-emerged, took a mean of 12 days to receive confirmatory DFAT results (Formstone et al. 2026), and sequence data took several months to generate (Yuson et al. 2024). Such delays limit the usefulness of genomic data for immediate outbreak response.

Decentralised sequencing capacity has expanded in many settings but remains uneven and is often concentrated around higher-priority pathogens and well-supported laboratories (Brunker et al. 2020; Li et al. 2022; Bacus et al. 2021; Lushasi et al. 2023; Jaswant et al. 2024; Gigante et al. 2020; Tan et al. 2026; Brito et al. 2021). Expanding sequencing capacity is worthwhile, but will take time, because sequencing is a highly skilled task that requires specialized infrastructure. For diseases such as rabies, where case numbers are lower and surveillance systems often more fragmented, dedicated sequencing capacity at every regional laboratory may be neither feasible nor necessary. Whether sequencing remains centralised or becomes increasingly decentralised, practical approaches to stabilising and transporting samples will remain essential. Long transit times, unreliable cold-chain infrastructure, high ambient temperatures and limited storage capacity, all common issues in rabies-endemic regions, can compromise RNA integrity and delay analysis. Specimen types that are stable at ambient temperature and fit within existing diagnostic workflows are therefore particularly attractive, as they avoid additional collection procedures and dedicated cold-chain transport.

Two candidate specimen types are Flinders Technology Associates (FTA) cards and lateral flow devices (LFDs). FTA cards preserve nucleic acids and inactivate pathogens, enabling safer transport without continuous cold-chain conditions. They have supported RABV detection, and in some settings, whole-genome sequencing from brain tissue (Wacharapluesadee et al. 2003; Picard-Meyer et al. 2007; Goharriz et al. 2017; Zhang et al. 2025; Sun et al. 2026). Their performance across a broader range of field-relevant storage and transport conditions remains unclear. LFDs are already used in many settings to support decentralised rabies diagnosis, with improved case detection and more timely initiation of post-exposure prophylaxis (Saito et al. 2025; Orciari et al. 2026; Naïssengar et al. 2021). Therefore, residual material on the device could provide genomic material without preparing and transporting an additional specimen. RABV RNA has been detected from LFDs after prolonged storage at ambient temperature (up to 83 days; Mauhay et al. 2023), and LFD-derived nucleic acids have supported whole-genome sequencing of other viruses, including SARS-CoV-2, dengue, and monkeypox (Pronier et al. 2026; Rector et al. 2023; Martin et al. 2022). However, PCR detection requires only short target regions, whereas whole genome sequencing requires sufficient RNA quality and quantity across the genome. Whether LFDs can reliably support RABV whole-genome sequencing under routine surveillance conditions has not been established.

Here, we evaluate FTA cards and LFDs as alternative specimen types for rabies genomic surveillance, in the context of the Philippines, where geographical isolation and complicated transportation pose logistics challenges. We first assess the diagnostic sensitivity and specificity of the BioNote LFD against DFAT under field conditions, then evaluate RABV RNA preservation and genome recovery from FTA cards and LFDs under controlled, field-relevant storage conditions, and from LFDs collected during routine surveillance in the MIMAROPA region. By linking field diagnostic validation with subsequent genomic analysis, we assess whether these specimens offer a practical route to reduce delays in specimen transport and sequencing in both centralised and decentralised rabies surveillance systems.

## Materials and Methods

### Study context and samples

The study comprised two complementary components: laboratory-based experiments to evaluate RNA preservation and genome recovery from LFDs under controlled conditions, and field-based evaluations to assess LFD diagnostic performance and the feasibility of recovering RABV genomes from LFDs collected during routine surveillance in the MIMAROPA region (also known as the Southwestern Tagalog Region, Region IV-B), of the Philippines.

### Ethics

The study was assigned RITM Research Accession Number 2019-0038, and the animal procedures were reviewed and approved by the RITM Institutional Animal Care and Use Committee (IACUC) under IACUC Certification no. 2019-001.

### Study site

MIMAROPA is an island region in the Philippines comprising the provinces of Mindoro (Oriental and Occidental), Marinduque, Romblon and Palawan (Figure 1 and Supplementary Figure 1). Although Romblon and Marinduque had previously been declared rabies-free, rabies re-emerged in Romblon in October 2022, with the first detected cases on the island of Tablas, followed by re-emergence in Marinduque in May 2023. Since 2020 Integrated Bite Case Management (IBCM) has been introduced to selected provinces in MIMAROPA to enhance rabies surveillance, as part of the SPEEDIER implementation research project (Yuson et al. 2024; Formstone et al. 2026), with 319 cases confirmed during the period covered by this analysis (August 2022 to July 2026). IBCM involves sample collection and LFD testing at the provincial level, followed by submission to Regional Animal Disease Diagnostic Laboratories (RADDLs) and/or the Rabies National Rabies Laboratory in the Research Institute for Tropical Medicine (RITM) in Manila for confirmatory testing by DFAT and sequencing.

**Figure 1.**
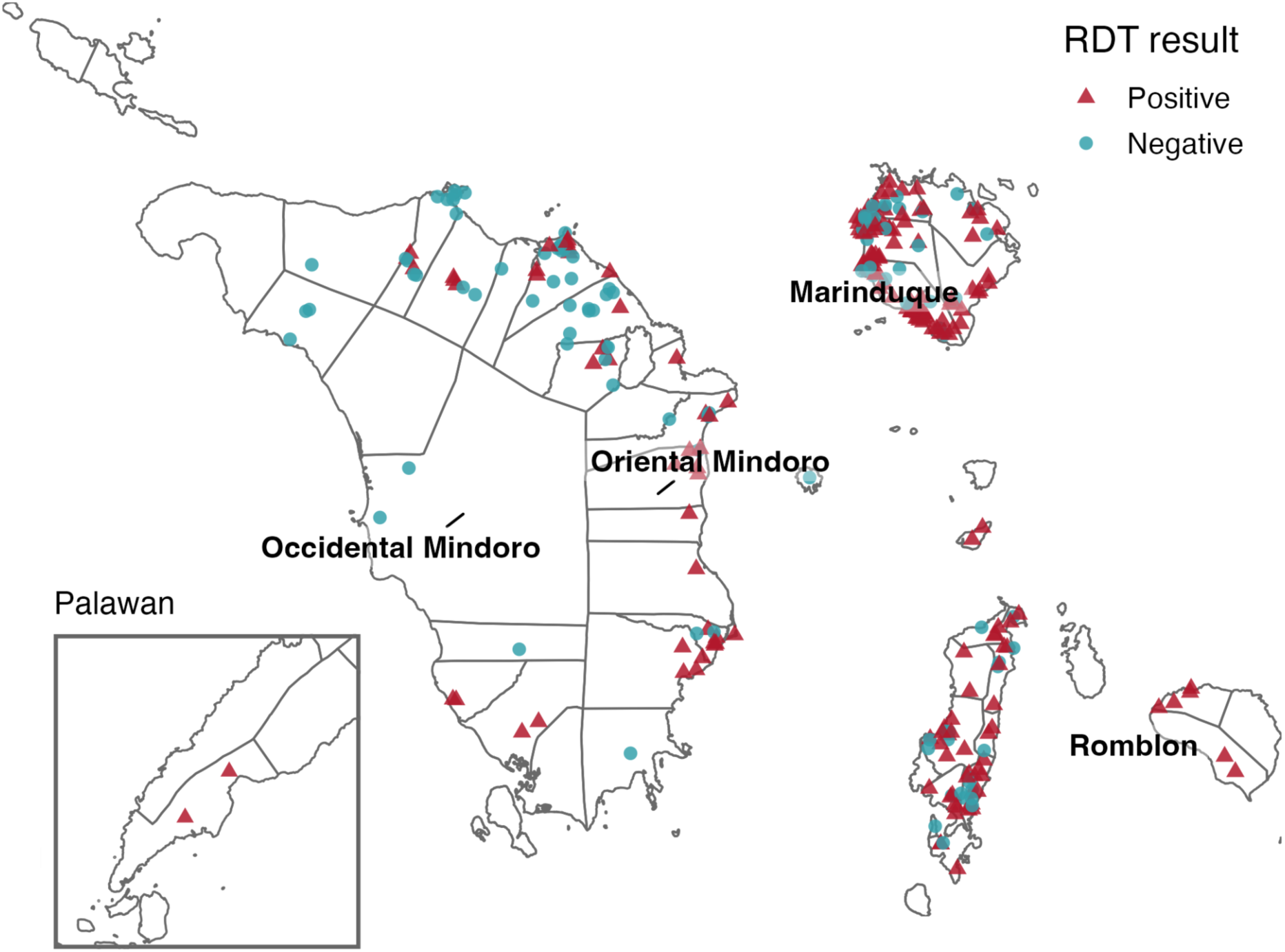
Locations of samples and the results of Lateral Flow Devices (LFDs) in MIMAROPA region, Philippines. Municipal boundaries are shown in grey, and province names are labelled. Since Palawan spans a large geographic extent and LFDs were performed within a relatively small area of the island, an enlarged inset shows the Palawan LFD survey area.

### LFD validation

Animal brain samples (n=319) for rabies testing in MIMAROPA were collected using the straw method through the occipital foramen (Barrat and Blancou 1988). These were tested using Bionote LFDs (Anigen Rapid Rabies Antigen Test, Bionote) in field and laboratory settings, following the manufacturer’s instructions, by trained animal health workers and/or licensed veterinarians. For Oriental Mindoro all LFDs were conducted at RADDL concurrently with DFAT; some samples were stored for a period of time before testing. In the other provinces, samples were tested at the point of collection and stored by the relevant Provincial Veterinary Office (PVO) and submitted to RADDL and/or RITM for confirmatory DFAT. Of 319 samples, 190 (59.6%) were tested in the field immediately after collection using the LFD, while 129 (40.4%) were LFD tested at the RADDL at the same time as DFAT testing. Specimens with a high epidemiological or clinical suspicion of rabies that were unsuitable for DFAT due to their condition were referred to RITM for confirmatory RT-qPCR.

### Laboratory turnaround times

Turnaround times (TATs) were calculated separately for diagnostic and sequencing workflows for cases from the MIMAROPA region included in the LFD validation. Diagnostic TATs were calculated for all cases with a recorded sample collection date, with stage-specific TATs calculated between sample collection, testing by LFD, and by confirmatory DFAT. Sequencing TATs were calculated for cases with a recorded sequencing date, with overall TAT defined as the number of days between confirmatory DFAT and sequencing.

### Samples for sequencing

Twenty rabies-positive dog brain tissues archived in 2025 and 2026 were obtained from the Rabies National Rabies Laboratory biobank at RITM (Supplementary Table 1). These samples were previously confirmed positive for rabies using DFAT. The frozen brain samples, which were stored without a preservation buffer, were thawed on ice before application to FTA cards and LFDs. As a negative control, an archived negative sample was also included. Additionally, eight rabies-positive LFD samples collected during initial field screening in MIMAROPA were included to represent field specimens stored at ambient temperature for 1-77 days before shipment to the RITM. These were stored at −20°C upon receipt then extracted after 6 days.

### Preservation experiments

Brain tissue samples were preserved on both FTA cards (Qiagen QIAcard FTA Classic Cards, Qiagen) and LFDs (Anigen Rapid Rabies Antigen Test, Bionote) to evaluate RNA stability under field-relevant conditions. Henceforth, we use ‘biological sample’ for the original brain tissue from an animal, and ‘specimen’ for each individual preparation of a biological sample that underwent RNA extraction – whether as an FTA card or LFD preparation under a given storage condition, or as a fresh-tissue aliquot processed directly at day 0. For FTA cards, a pea-sized amount (∼125 mm^3^) of brain tissue was applied to each sample spot using sterile swabs, ensuring complete absorption into the matrix. LFDs were used according to the manufacturer’s instructions. Following application, both FTA cards and LFDs were air-dried for 4 hours in a biosafety cabinet, individually sealed in resealable bags containing 3g of desiccant, and placed in an outer resealable bag and storage box to minimise moisture and direct sunlight exposure.

In the main experiment (cohort 1), five samples were evaluated across eight storage conditions defined by temperature (ambient [25–30°C] or 37°C) and time post-preservation (1, 3, 7, and 21 days), with RNA extracted at each condition. A follow-up experiment (cohort 2) with an additional 15 samples assessed storage at ambient temperature only, with RNA extraction at days 3 and 7. RNA extracted from fresh brain tissue on day zero was processed in parallel as a baseline comparator. The full set of conditions is shown in Table 1. A rabies-negative brain sample was processed alongside study samples as a negative extraction control. In cohort 1, additional negative controls were also prepared by applying water, rather than brain tissue, to both LFDs and FTA cards.

**Table 1.**
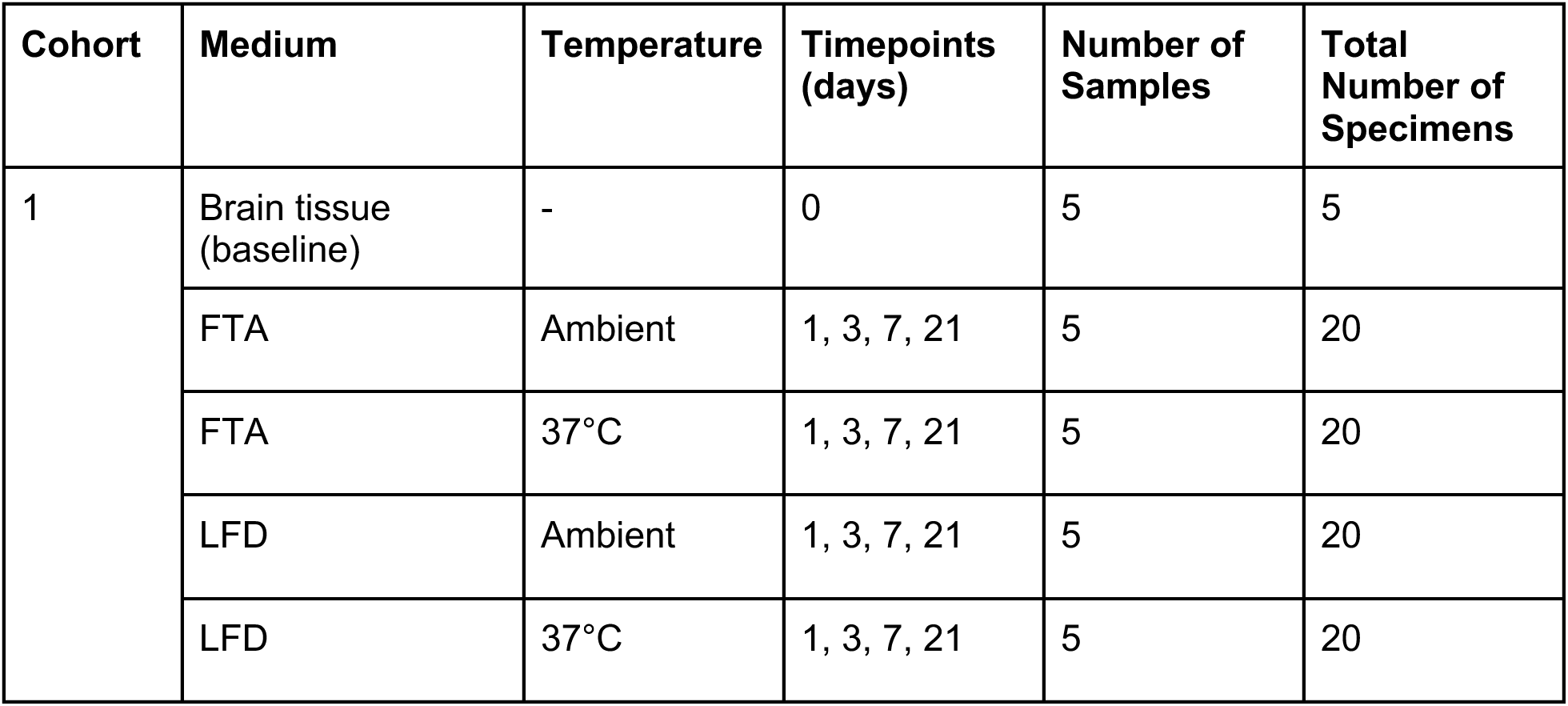
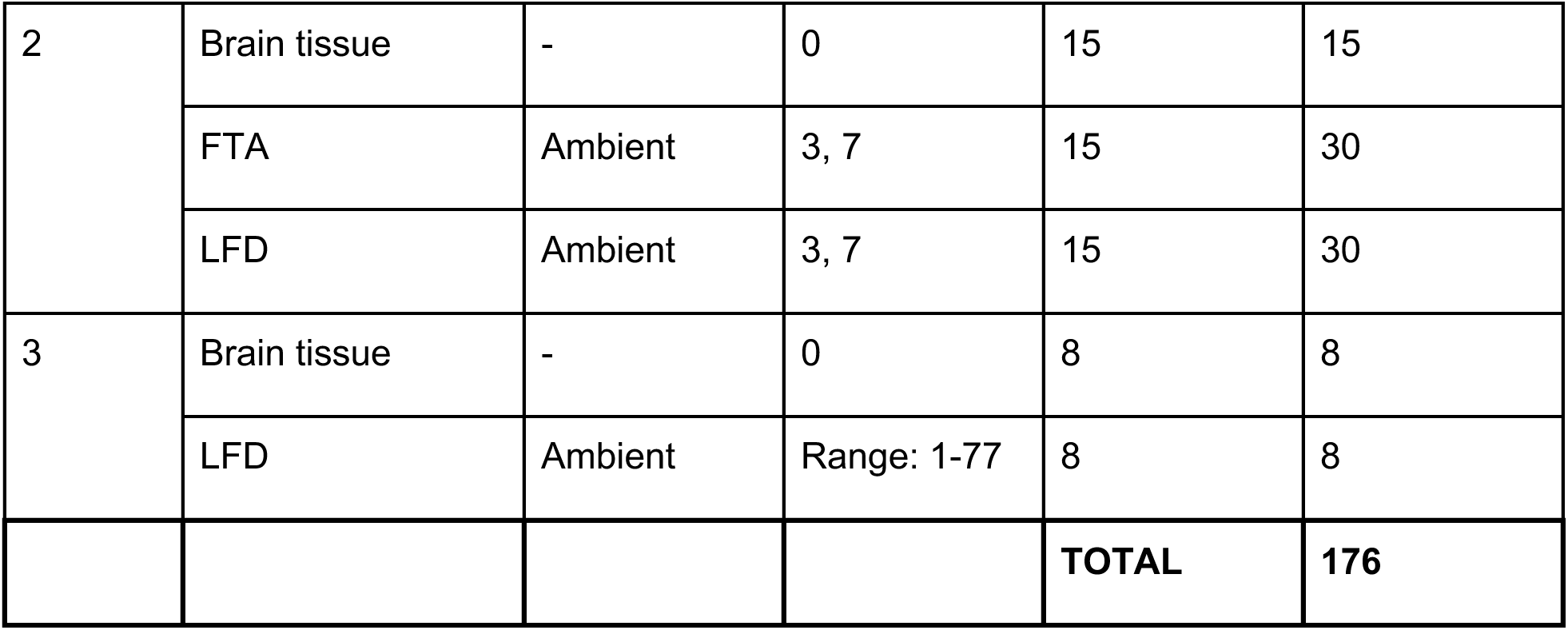
Study design and distribution of rabies virus specimens across experimental cohorts, storage conditions and timepoints. For each biological sample, the baseline brain-tissue specimen and matched FTA-card and LFD specimens were prepared from aliquots of the same original brain-tissue sample. Cohort 1 evaluated lateral flow devices and FTA-card specimens under controlled storage conditions at ambient temperature and 37°C for up to 21 days. Cohort 2 evaluated specimens stored at ambient temperature for 3 or 7 days, and Cohort 3 comprised specimens collected and stored under routine field conditions.

| Cohort | Medium | Temperature | Timepoints (days) | Number of Samples | Total Number of Specimens |
| --- | --- | --- | --- | --- | --- |
| 1 | Brain tissue (baseline) | - | 0 | 5 | 5 |
|  | FTA | Ambient | 1, 3, 7, 21 | 5 | 20 |
|  | FTA | 37°C | 1, 3, 7, 21 | 5 | 20 |
|  | LFD | Ambient | 1, 3, 7, 21 | 5 | 20 |
|  | LFD | 37°C | 1, 3, 7, 21 | 5 | 20 |
| 2 | Brain tissue | - | 0 | 15 | 15 |
|  | FTA | Ambient | 3, 7 | 15 | 30 |
|  | LFD | Ambient | 3, 7 | 15 | 30 |
| 3 | Brain tissue | - | 0 | 8 | 8 |
|  | LFD | Ambient | Range: 1-77 | 8 | 8 |
|  |  |  |  | **TOTAL** | **176** |

### RNA extraction

From each FTA card sample spot, an 80-100 mm² square was excised using a sterile blade. For LFDs, the sample pad was divided lengthwise into two strips using a sterile blade, and one half was used per extraction. FTA cards or LFD sample pads (half-strip per extraction) were cut into small pieces and incubated in 120 μL nuclease-free water for 30 min at ambient temperature with intermittent vortexing. Following centrifugation (maximum speed, 2 min), 110 μL of eluate was recovered and processed using the RNeasy^®^ Mini Kit (Qiagen) according to the manufacturer’s instructions, including on-column washes and elution in 60 μL nuclease-free water. RNA concentration was measured using the Qubit^TM^ RNA High Sensitivity Kit (Thermo Fisher Scientific). Extracted RNA was aliquoted and stored at −80°C. A detailed step-by-step protocol for LFD and FTA card preparation and extraction, including photographs, is available on protocols.io (Crisostomo et al. 2026).

### RT-qPCR

A real-time SYBR-based reverse transcription PCR (RT-PCR) assay was used to detect lyssavirus RNA in samples (Marston et al. 2019). RNA extracted from brain tissue, FTA cards, and LFDs was used as templates and β-actin was included as an internal control to assess RNA integrity and successful amplification. Positive and negative controls were included in each run. Changes in the quantification cycle (Cq) relative to baseline brain tissue were used as a proxy for RNA degradation under each condition. Changes in Cq values relative to baseline brain tissue were calculated as ΔCq = Cq_alternative specimen − Cq_baseline brain tissue. The effects of preservation medium (FTA card or LFD), storage temperature (ambient or 37°C), and storage duration (1, 3, 7, and 21 days) on ΔCq values were assessed using a linear mixed-effects model, with sample included as a random effect to account for repeated measurements from the same original brain sample.

### Nanopore sequencing

An established sample-to-sequence protocol was used to generate amplicon-based libraries for nanopore sequencing (Bautista et al. 2023). A multiplex PCR reaction was performed using Q5 High Fidelity Hot-Start DNA Polymerase (New England Biolabs, UK) and a Philippines RABV-specific multiplex primer set (Brunker et al. 2020). Library preparation was performed using a Nanopore Native Barcoding Kit 24 V14 (SQK-NBD114.24, Oxford Nanopore Technologies, UK). Negative controls were included in each run to monitor cross-contamination. The final library was loaded onto an R10 flow cell and sequenced on a MinION device with live basecalling. Reads were processed using an established bioinformatic pipeline (https://github.com/RAGE-toolkit/Artic-nf; Bautista et al. 2023).

### Analysis of sequencing results

To evaluate variation in genomic coverage across conditions, the percentage of the RABV genome achieving at least 20x read depth, non-masked coverage, was modelled using a beta mixed-effects regression framework. Analyses were restricted to FTA and LFD samples from cohort 1, with storage temperature (RT or 37°C), sample type (FTA or LFD), and timepoint (days 1, 3, 7 and 21) included as fixed effects. Sample identity was included as a random intercept to account for repeated measurements. A full three-way interaction was tested and removed when unsupported by a likelihood-ratio test. Model fit was assessed using DHARMa simulated residual diagnostics. Estimated marginal means and Tukey-adjusted pairwise comparisons were used to assess differences between sample types, temperatures and timepoints.

### Consensus sequence analysis

Lineage designation was performed on consensus sequences using MADDOG V2 (https://github.com/RAGE-toolkit/MADDOG-V2; Campbell et al. 2022). To assess concordance between consensus sequences generated from the same biological sample under different specimen types, storage conditions and timepoints, pairwise nucleotide distances were calculated between all available within-sample consensus sequence pairs. Comparisons were restricted to aligned positions with unambiguous nucleotide calls (A, C, G or T) in both sequences, excluding positions containing ambiguous or masked bases. Pairwise distances were expressed as the number of single nucleotide polymorphisms (SNPs) between each sequence pair. Associations between SNP differences and specimen type, storage temperature, storage duration, amplicon failure and read depth were assessed descriptively.

### Data and code availability

All data generated and analysed during this study, together with the code used for data processing, analysis and visualisation, are available in https://github.com/RAGE-toolkit/RABV-LFD-sequencing.

## Results

### LFD validation

A total of 319 samples had results for both LFD and DFAT tests available. Tests were conducted between August 2022 and July 2026 across five provinces in the MIMAROPA region: Oriental Mindoro (n = 87), Occidental Mindoro (n = 11), Marinduque (n = 146), Romblon (n = 73), and Palawan (n = 2), see Figure 1. The majority of the tests were conducted on dogs (n=276), followed by cats (n=28), pigs (n=7), cattle (n=3), goats (n=3), and sheep (n=2). There was one discordant LFD result (negative test conducted by PVO, positive test conducted by RADDL); with the RADDL test result retained for analysis. Using the DFAT as the gold standard, LFD sensitivity was 99.1% (210/212; 95% CI 96.6-99.7%) and specificity was 100% (107/107; 95% CI 96.6-100.0%) (Table 2).

**Table 2.** LFD performance against DFAT. The DFAT was considered gold standard, N = 319

|  | DFAT Positive | DFAT Negative |
| --- | --- | --- |
| LFD Positive | 210 | 0 |
| LFD Negative | 2 | 107 |

### RT-qPCR detection of rabies virus RNA

RABV RNA was detectable by RT-qPCR in 169 of 176 specimens compared in the experimental cohorts (Table 1), with quantification cycle (Cq) values ranging from 14.16 to 35.39 (Supplementary Table 2). In cohort 1, sample identity accounted for a substantial proportion of the variability in ΔCq (ICC = 0.84). Preservation medium had a strong effect on ΔCq (F_1,58_ = 320.36, p < 0.001): LFD samples had a mean ΔCq 5.00 cycles higher than FTA samples (SE = 0.28, p < 0.001), indicating a substantially greater reduction in detectable viral RNA signal in LFD-preserved samples. Storage at 37°C was associated with a small increase in ΔCq compared with ambient temperature (0.58 cycles, 95% CI 0.02-1.14; F_1,58_ = 4.34, p = 0.042). Storage duration also influenced ΔCq (F_3,58_ = 7.93, p < 0.001), although differences were modest, with ΔCq at day 21 approximately 1.2-1.8 cycles higher than at earlier timepoints. No significant interactions between preservation medium, temperature, and storage duration were observed.

Contamination was detected during day 3 processing of cohort 1 FTA-card samples, with RABV RNA amplified from the negative extraction control (a rabies-negative brain sample) and β-actin RNA detected in the water-only control. Due to limited equipment access, RT-qPCR screening was performed either before or after sequencing, with RNA extracts frozen pending testing when screening could not be performed immediately. Consequently, contamination in this batch was identified retrospectively, after sequencing.

### Sequencing results

#### Overall performance and genome recovery

A total of 176 specimens (brain, FTA card & LFD from the cohort experiments) were sequenced across 9 runs. One run was found to have contamination in the negative template control and was excluded from the analysis, resulting in the removal of three cohort 2 samples (N6,10,11). Of the remaining 161 specimens, one day-1 FTA card produced no mapped reads and was excluded as a likely extraction failure, leaving 160 specimens for analysis. Full specimen-level details for all 160 specimens are provided in Supplementary Table 2.

Overall, 131/160 (81.9%) achieved ≥90% genome coverage at ≥20x read depth. Genome recovery differed by specimen type, with the highest genome recovery from baseline brain specimens (96.0%), followed by FTA cards (85.7%) and LFD (73.6%) (Figure 2). Across the laboratory cohorts, FTA-card and LFD specimens generally achieved higher genome recovery than corresponding baseline brain specimens, with the largest difference observed in cohort 1, where none of the archived brain specimens reached ≥90% coverage (Figure 2).

**Figure 2.**
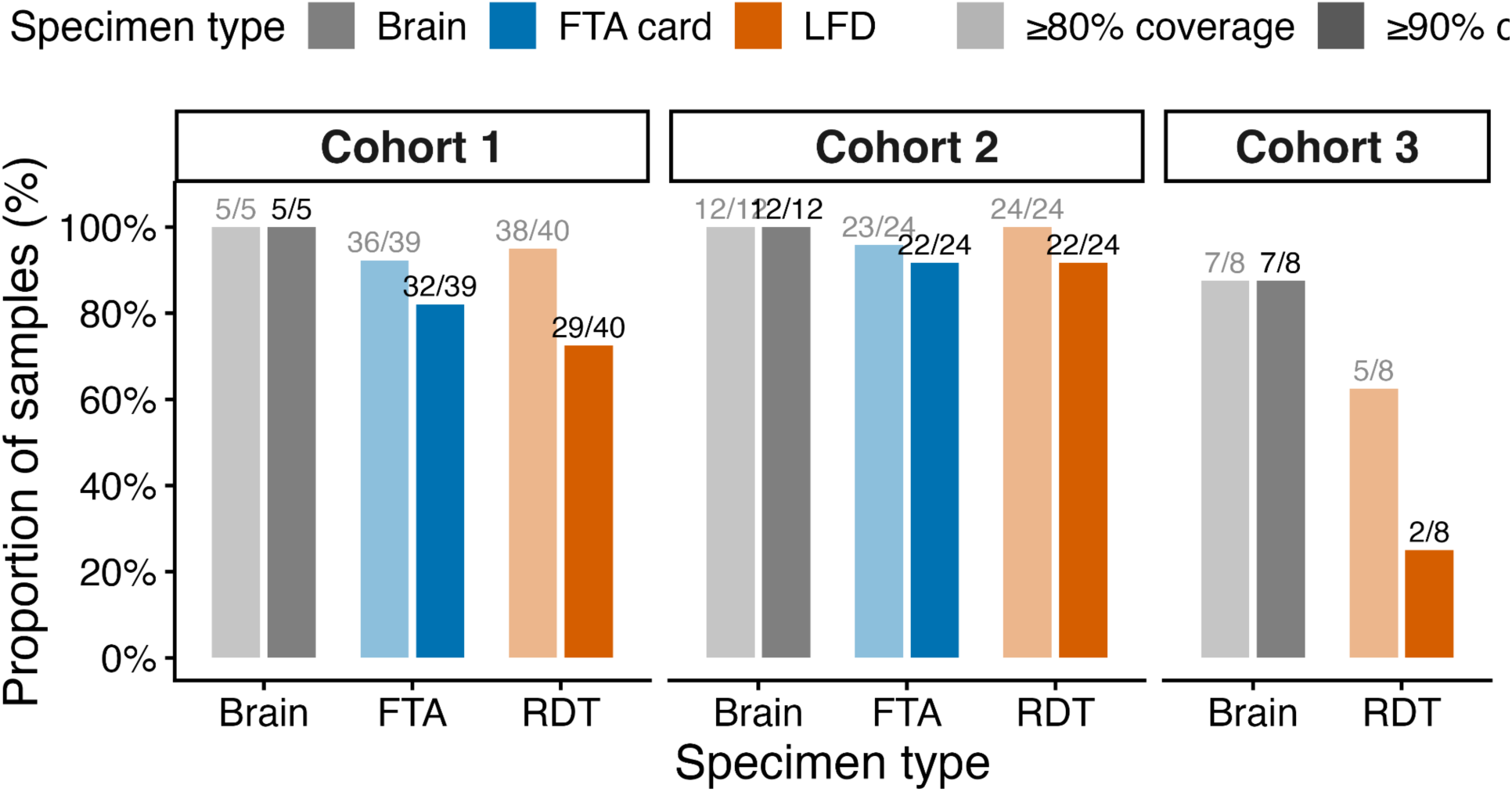
Genome recovery by specimen type and sequencing cohort. Proportion of specimens achieving ≥80% and ≥90% genome coverage at ≥20x read depth, shown by specimen type across the three cohorts. Bars are coloured by specimen type (brain, grey; FTA card, blue; lateral flow device [LFD], orange), with lighter shading indicating the ≥80% threshold and darker shading indicating the ≥90% threshold. Cohort 3 did not include FTA-card specimens.

Genome recovery was particularly high in cohort 2, with 91.7% of FTA-card and 91.7% of LFD specimens achieving the threshold 90% coverage following storage for 3 to 7 days at ambient temperature. In the field validation cohort (cohort 3), recovery was more variable: 2/8 (37.5%) LFD and 7/8 (87.5%) brain specimens achieved ≥90% coverage.

#### Effects of storage conditions on genome recovery

In cohort 1, which assessed performance across the full range of storage conditions (Figure 3), 61/79 (77.2%) of FTA-card and LFD specimens achieved ≥90% genome coverage, and 74/79 (93.7%) achieved ≥80% coverage (Figure 2). Of the five specimens with <80% coverage, four had been stored for 21 days and four were derived from biological sample 4. All cohort 1 brain tissue samples reached >96% genome coverage.

**Figure 3.**
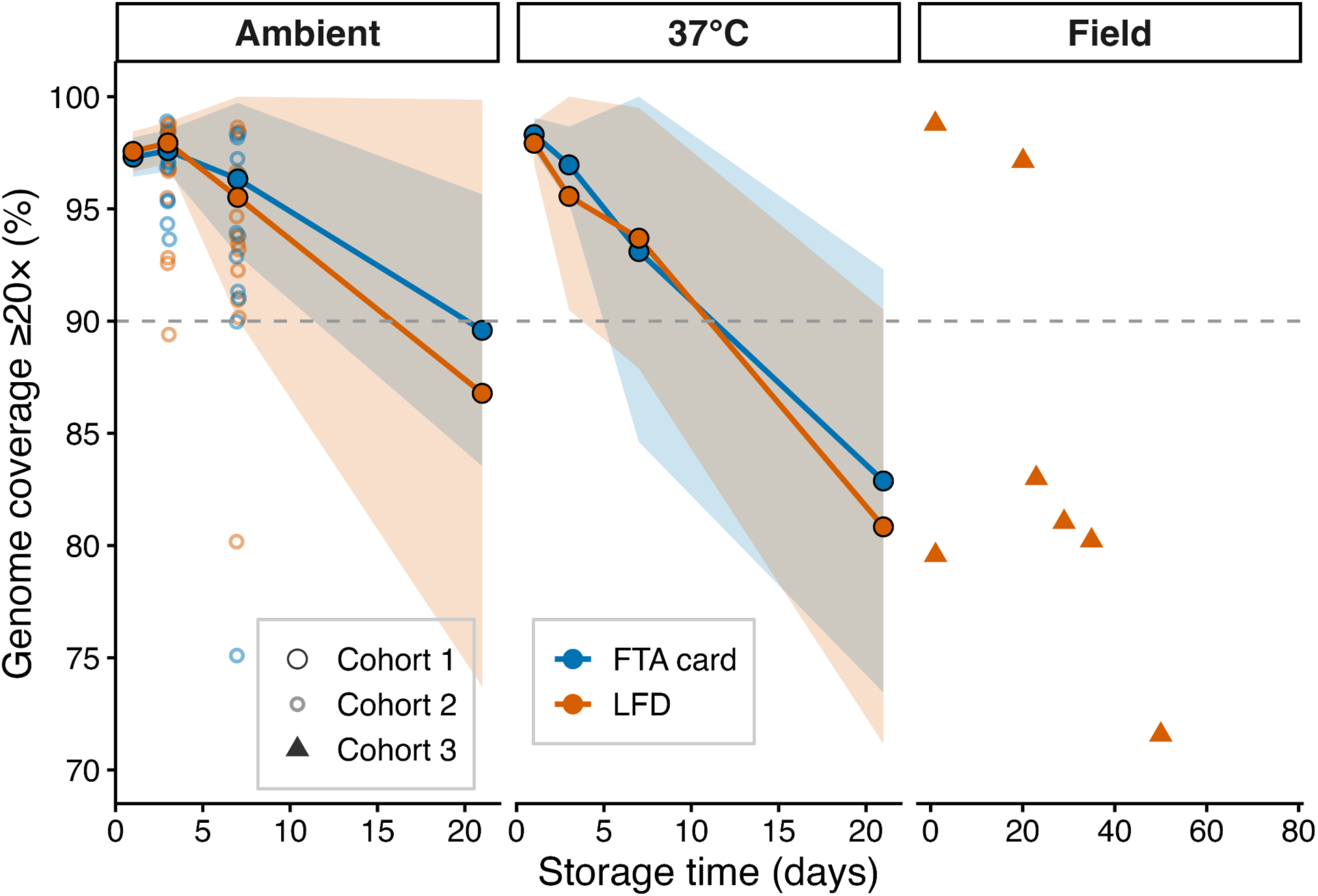
Rabies virus genome recovery from FTA card and lateral flow device (LFD) specimens over time. Percentage genome coverage at ≥20x depth (%) following storage of rabies virus-positive specimens under ambient temperature and 37°C conditions. Cohort 1 shows mean genome coverage ± standard deviation for FTA-cards and LFDs across storage time points. Cohort 2 shows individual validation samples, and Cohort 3 shows individual field-collected samples. Colours indicate preservation method (FTA card or LFD), while point shapes indicate cohort. The dashed horizontal line indicates 90% genome coverage.

Genome coverage declined with prolonged storage, with differences according to storage temperature (Figure 3). A beta mixed-effects model identified a significant association between storage duration and coverage (Type III Wald χ²₃= 56.00, p < 0.001), but no evidence of an overall specimen-type effect or interactions involving specimen type. Coverage was lower at 37 °C than at ambient temperature at days 7 (OR for ambient temperature vs 37 °C = 1.59, 95% CI 1.08–2.35, p = 0.019) and 21 (OR = 1.74, 95% CI 1.33-2.26, p < 0.001), but not at days 1 or 3. Variation between biological samples was substantial (random-intercept SD = 0.54, logit scale).

Coverage did not differ significantly between LFD and FTA-card specimens at any time point. At day 21, estimated coverage was 85.8% for LFD specimens and 88.4% for FTA-card specimens (OR for FTA vs LFD = 1.25, 95% CI 0.96-1.63, p = 0.099). Coverage declined over time in both specimen types, with significantly lower coverage at day 7 than day 1 for both FTA cards (p = 0.004) and LFDs (p = 0.008), and further declines by day 21 compared with earlier time points (p < 0.001).

#### Amplicon failure

Rates of amplicon failure were rare, with only 1 out 41 amplicons (amplicon 41) showing a failure rate exceeding 20% (Supplementary Figure 2A). Within this overall low range of failure rates, failure tended to increase over time, particularly by day 21, with a more pronounced increase in LFD samples than in FTA cards (Supplementary Figure 3). The highest failure rates were observed in LFD samples at day 21 and in baseline brain tissue at day 0. The higher apparent amplicon failure in baseline brain tissue coincided with an increase in longer-than-expected RABV reads, potentially reflecting better-preserved RNA that supported amplification of longer products rather than the expected ∼400 bp targets. This may have contributed to the apparent amplicon failure without necessarily indicating poorer sample quality.

#### Genome consensus variation

For the 160 specimens included in downstream analysis, there were 792 within-sample pairwise comparisons (successful sequencing runs across 25 samples, according to 1-3 timepoints and 1-3 storage conditions, Table 1). All consensus sequences derived from the same biological sample received the same lineage assignment, irrespective of specimen type, storage condition or timepoint. The median pairwise distance was one SNP (IQR 0-1; range 0-5), with 47.1% of comparisons showing identical consensus sequences. The highest SNP count was five, observed in a day 3 FTA-card sample (S5 in Figure 4) from the batch in which contamination was detected by RT-qPCR. There was no clear association between the number of SNPs and specimen type, temperature, storage duration (data not shown). Similarly, there was no association with amplicon failure and SNP position (Supplementary Figure 2b), or read depth and SNP position (Supplementary Figure 4). Instead, variation appeared to be primarily sample-specific, with some biological samples showing consistently greater differences between their corresponding consensus sequences than others, although levels of variation were also higher in the cohorts subjected to controlled laboratory storage conditions (cohorts 1 and 2; Figure 4). The highest concordance was observed among the field-derived samples processed in Cohort 3. Of eight paired LFD- and brain-derived consensus sequences, seven were identical and the remaining one differed by a single SNP. Samples had been stored for 1-77 days prior to sequencing, with the SNP difference occurring in a sample stored for 29 days.

**Figure 4.**
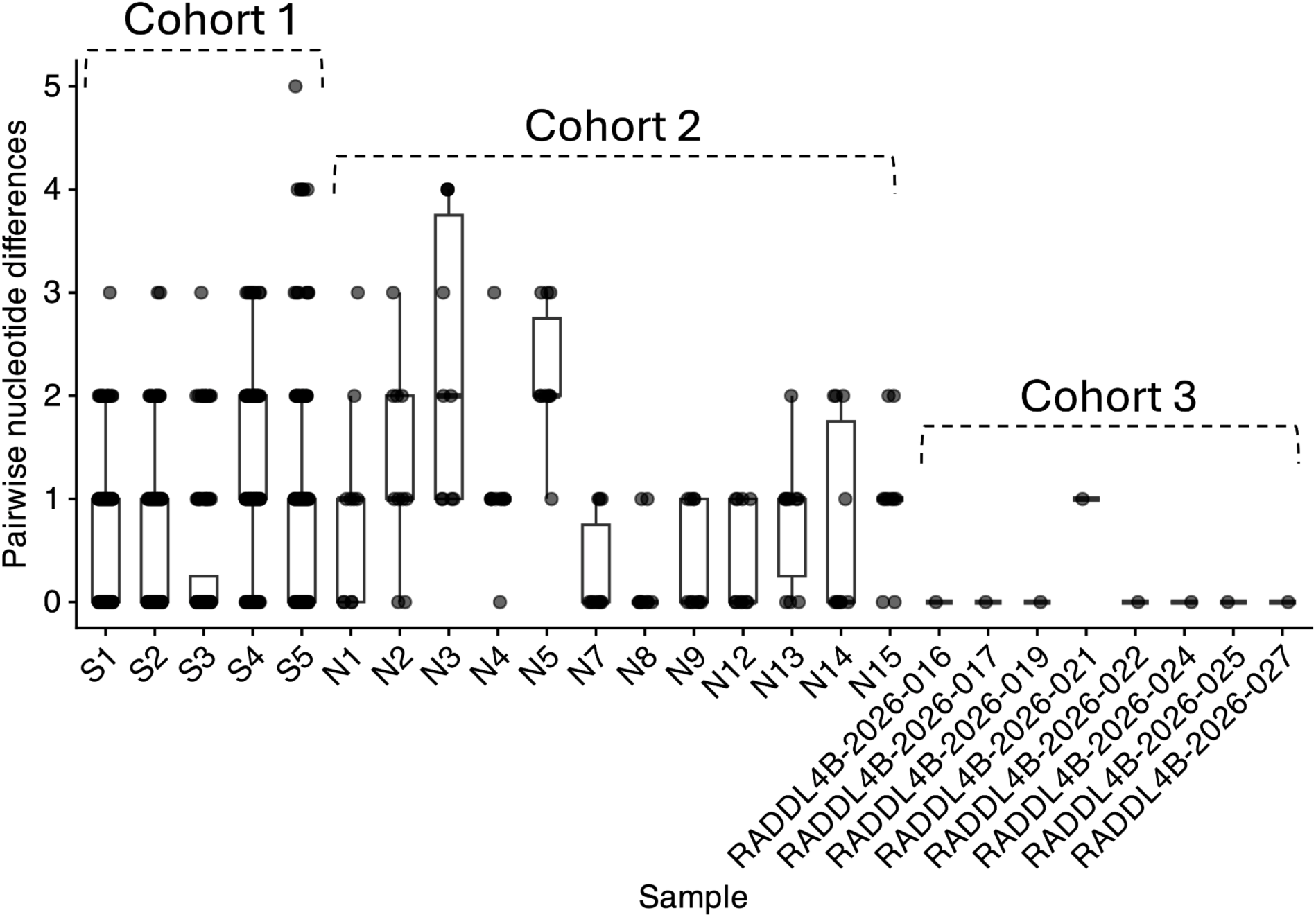
Pairwise genetic distances between consensus sequences. Consensus sequences were generated from the same biological samples (n=25) across different specimen types (brain, FTA card, LFD, and storage duration).

#### Turnaround times

Out of the 319 cases tested by LFD and DFAT in MIMAROPA, LFD was performed a median of 1 day (Interquartile range (IQR), 3 days) after sample collection, with confirmatory DFAT performed a median of 5 days (IQR, 12.2 days) later. The calculation included 300 cases with complete dates for both sample collection and LFD testing, and 308 cases with complete LFD and DFAT dates.

Among sequenced cases (Figure 5), the median time from DFAT confirmation to sequencing was 321 days (IQR, 167 days), with a significant reduction in turnaround time over the study period (linear regression: p < 0.001, R^2^ = 0.73). This reduction coincided with a shift towards smaller sequencing batches and shorter intervals between confirmatory dFAT and sequencing later in the study. The two cases sequenced in 2026 had a sequencing turnaround of less than 60 days.

**Figure 5.**
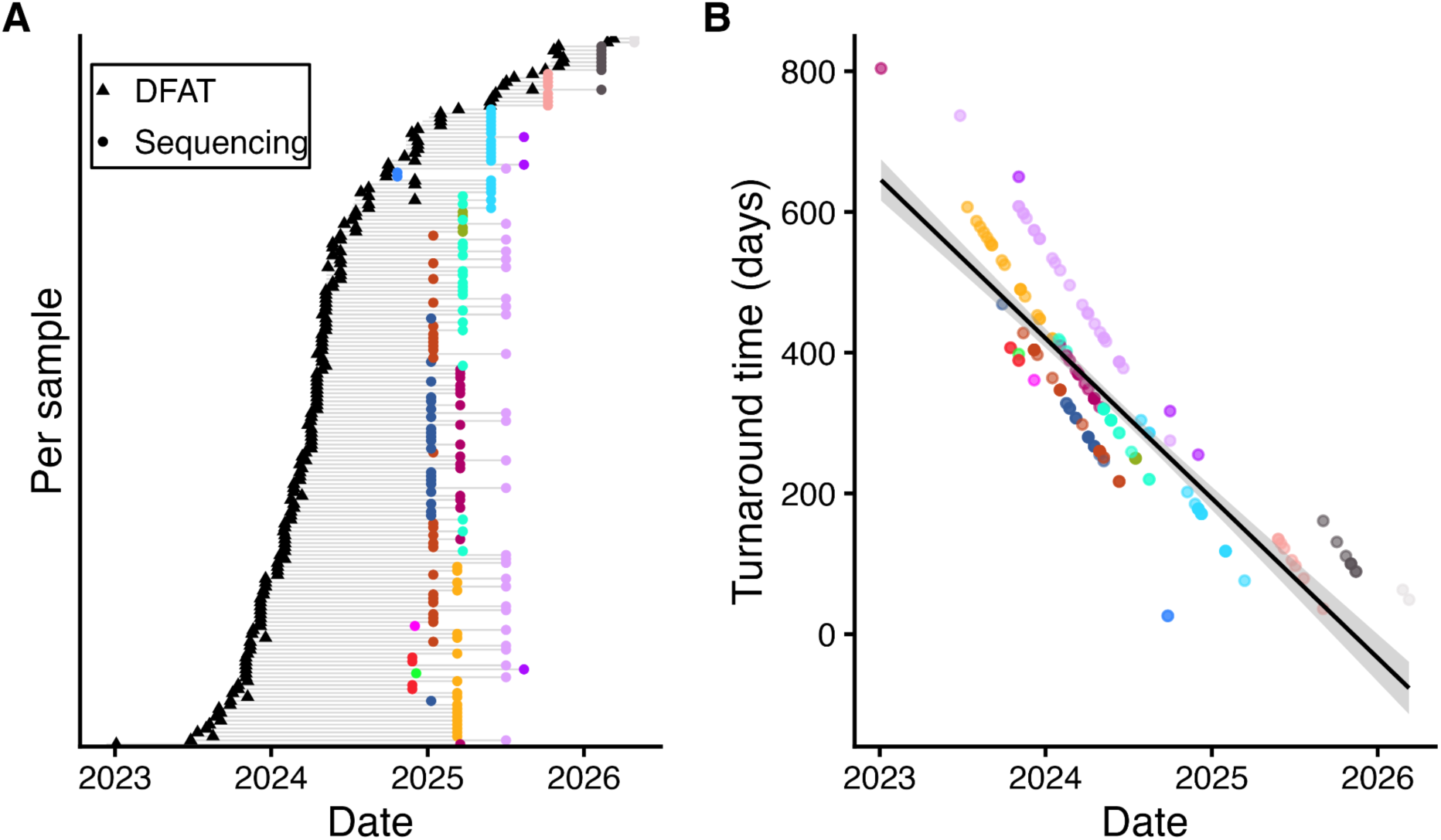
Turnaround time from rabies direct fluorescent antibody test (DFAT) confirmation to sequencing. (A) Individual sample timelines from sample collection to sequencing. Triangles indicate confirmatory DFAT testing and circles indicate sequencing; colours denote sequencing runs. (B) Time from DFAT confirmation to sequencing plotted against the date of DFAT confirmation. The solid line represents the linear regression fit, with the shaded area indicating the 95% confidence interval.

## Discussion

Rabies genomic surveillance is constrained not only by access to sequencing infrastructure and capacity, but by the time, cost, and logistical effort of obtaining samples from dispersed collection sites and transporting them safely to a sequencing facility. We show that BioNote LFDs already used for frontline rabies diagnosis can provide residual material suitable for whole-genome sequencing, linking rapid diagnosis to genomic surveillance without a separate specimen being collected, preserved, and transported. LFDs showed high concordance with DFAT across multiple host species, and LFD-derived material supported recovery of near-complete RABV genomes under both laboratory-simulated field conditions and routine field surveillance.

The BioNote LFD had a sensitivity of 99.1% and specificity of 100% during an active outbreak investigation in the Philippines, where it was used operationally by frontline personnel. This is consistent with previous evaluations of BioNote and Adtec rabies LFDs in the Philippines (sensitivity 94-96%, specificity 99.7-100%; Kimitsuki et al. 2020; Mananggit et al. 2021; Cruz et al. 2023) and elsewhere (sensitivity 92-97%, specificity 93-100%; Orciari et al. 2026; Tenzin et al. 2020; Sharma et al. 2015; Léchenne et al. 2016). These findings support BioNote LFDs as a robust frontline screening tool for rabies. The high sensitivity observed may partly reflect testing of fresh tissue shortly after collection, limiting antigen degradation (Orciari et al. 2026) both false negatives were generated by new users, consistent with an effect of user proficiency (Cruz et al. 2023). Training and competency assessment are therefore essential and a negative result should not be considered definitive when suspicion of rabies remains high– such specimens should be referred for confirmatory testing. However, LFD performance varies substantially across commercial products (Eggerbauer et al. 2016; Orciari et al. 2026), hence our findings are specific to BioNote and other brands will require equivalent evaluation.

In the laboratory cohorts, both FTA cards and LFDs supported near-complete genome recovery, although recovery declined after extended storage, particularly for LFDs and at higher temperatures. In the field cohort, near-complete genomes were also recovered from LFDs stored at ambient temperature for up to 23 days, whereas coverage declined progressively with longer storage, indicating a useful but limited window of around three weeks. These results were obtained with an established, widely used amplicon-based nanopore protocol (Brunker et al. 2020; Bautista et al. 2023; Quick et al. 2017) whose short (∼400 bp) amplicons tolerate degraded RNA and whose primer scheme can be adapted to other regions, supporting integration into existing sequencing workflows. Consensus sequences from the same biological sample were highly concordant (median 1 SNP), with differences mainly sample-specific and not linked to specimen type, storage, amplicon failure or read depth. This finding points to sample-specific factors, such as mixed-base calls from low-frequency intra-host variation at modest depth (Grubaugh et al. 2019; Liu et al. 2022), rather than systematic artefacts of LFD or FTA storage. Field LFD and brain sequences were identical in seven of eight pairs, while the maximum was 5 SNPs in an FTA-card sample. Differences of this size are unlikely to affect lineage assignment (Paull et al. 2024) but should be considered in fine-scale transmission inference.

The highest genome coverage was achieved from baseline brain-tissue samples extracted on day zero, as expected given their lower Cq values relative to other specimens (Supplementary Table 2). These samples also yielded longer reads, with read lengths shifting towards the expected 400 bp amplicon length over time and in FTA-card and LFD samples. This pattern likely reflects differences in RNA integrity: better-preserved RNA in baseline extracts may support amplification of longer RABV fragments, whereas RNA degradation may favour the recovery of shorter products. RNA degradation in brain tissue during storage would also be expected, particularly where refrigeration and nucleic acid preservation buffers are unavailable, as may be the case in resource-limited field settings. Although this was not directly assessed in the present study, it highlights the potential value of FTA cards and LFDs as alternatives for preserving RABV RNA when freezer storage is unavailable. The protocol was designed to maximise recovery from highly degraded samples (Brunker et al. 2020; Quick et al. 2017), reflecting the suboptimal storage and transport conditions to which field-collected RABV specimens may be exposed. The recovery of longer-than-expected products from higher-quality samples is therefore not unexpected, but may reflect the greater integrity of the available RNA.

FTA cards and LFDs have different operational roles. FTA cards are well suited to preserving samples for long-distance transport (Zhang et al. 2025) but require an additional workflow of sample application, drying, packaging and later card punching. This adds training, staff time, and consumables in settings already affected by staff and resource constraints (Formstone et al. 2026), and introduces additional biosafety and contamination risks. LFDs, by contrast, retain residual material within a single-use device already used in frontline screening in many settings. Once the result is read, the device can be stored and submitted for sequencing without a separate specimen or waiting for confirmatory testing and cold-chain shipment of the primary sample. This approach could reduce the marginal cost, handling burden, and delay associated with obtaining specimens for genomic surveillance. Where virus inactivation by the assay buffer has been validated, LFDs may offer an additional biosafety advantage (reported for BioNote by Eggerbauer et al. (2016), but not stated in the manufacturer’s documentation).

Realising these benefits depends on the surveillance system around the lateral flow device. LFD sampling removes a specimen bottleneck but will not by itself shorten sequencing turnaround. In MIMAROPA, turnaround improved as workflows were established, dedicated staff were assigned and smaller batches were processed, but delays in sample shipment, competing staff responsibilities, batch-based sequencing and limited analytic capacity remained. Sustainable implementation will require protected staff capacity, clear specimen-referral pathways, and sequencing arrangements that support timely analysis of small numbers of samples, for example, through combined-pathogen runs. A hub-and-spoke model (Saha et al. 2024; Kanteh et al. 2022), in which regional laboratories undertake local sample processing and targeted sequencing, particularly during outbreaks when speed matters most, while central laboratories provide quality assurance, specialised analyses, and data management, could be a pragmatic alternative to relying on few central sequencing laboratories. Combined with LFD sampling, such operational adaptations could shorten the path from diagnosis to genomic data to support action. LFD results themselves should also be used. Point-of-collection testing can incentivise frontline workers by providing an immediate result, and users have reported positively on their usefulness (Formstone et al. 2026; Lushasi et al. 2020). However, in many countries LFDs have no clear regulatory status and there is no national guidance on how to report or act on results. This uncertainty can delay the release of results and make personnel reluctant to act (Formstone et al. 2026). Realising the potential of LFDs will therefore require clear guidance on reporting and acting on their results, as well as validated workflows.

This study has several limitations. First, the field evaluation was small (n = 8): although five LFDs achieved ≥80% genome coverage, only two reached ≥90%. Storage duration alone did not explain variation in coverage, as one device stored for just one day achieved only 79.6% coverage. Larger field evaluations across host species, viral loads, storage conditions and settings are therefore needed to better characterise performance under routine surveillance conditions. Second, we did not optimise RNA extraction or sample preservation. Using the whole LFD strip rather than half may increase RNA recovery, while collecting brain tissue into a nucleic acid preservation buffer (e.g. DNA/RNA Shield) could improve RNA stability during storage, thawing and handling, although both approaches require further evaluation and may have cost implications. The laboratory comparisons also relied on archived brain tissue, and only the BioNote LFD was evaluated; performance may differ with other LFD brands, specimen types or workflows. Finally, contamination remains an important consideration when integrating molecular testing into routine surveillance. One sequencing run was excluded following contamination of the negative control, highlighting the need for robust quality-control procedures, including negative controls and physical separation of pre- and post-amplification activities. A signal in an extraction control from one batch of FTA-card extracts also raised the possibility of contamination during card cutting. Dedicated punchers, single-use cutting tools and processing smaller batches may help mitigate this risk.

Overall, this study shows that LFDs used for rabies diagnosis can double as genomic specimens: one test gives a diagnosis at the point of collection and, once read, a sample ready for sequencing, with no additional collection procedures and no cold chain requirements. Because LFDs may already be deployed where cases are first detected, this approach can bring peripheral sites into the genomic surveillance network, removing a major barrier to genomic surveillance: timely access to a suitable specimen. LFD-based diagnosis combined with genomic surveillance could strengthen case detection, support investigation of transmission and reintroductions, and broaden the geographic reach of rabies genomic surveillance. Use of LFDs does not replace confirmatory testing or robust laboratory systems, but offers a practical, potentially lower-cost route to genomic data where conventional specimen transport is slow, difficult or unaffordable.

## Acknowledgements

This work was funded by a UK Medical Research Council New 8Investigator Research Grant (MR/X002047/1) and Newton funding from the UK Medical Research Council (MR/R025649/1) and the8 Philippines Department of Science and Technology (DOST).

