## Supplementary Material for "Lateral flow devices can fast-track rabies virus genomic surveillance"

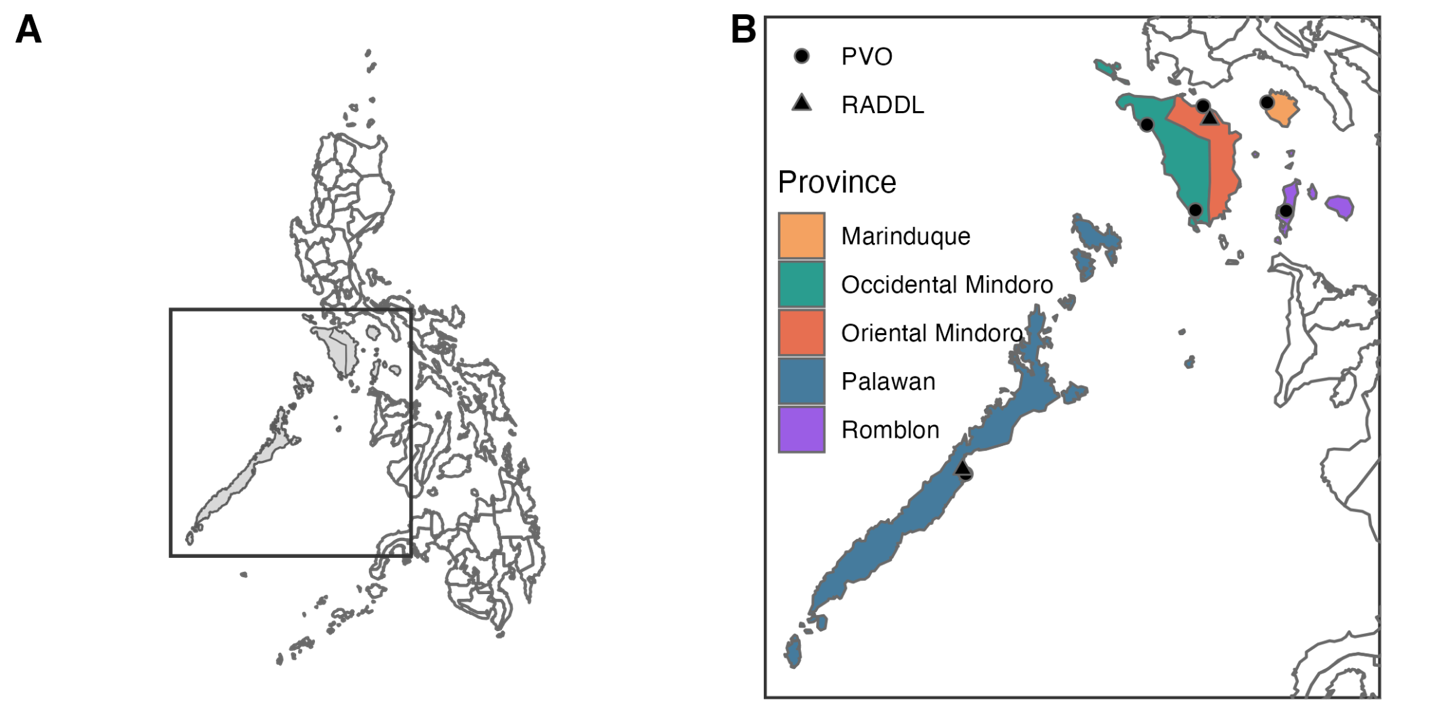

**Supplementary Figure 1. MIMAROPA region. (**A) Location of MIMAROPA within the Philippines; (B) The island provinces comprising the MIMAROPA region with locations of Provincial Veterinary Offices (PVOs) and Regional Animal Disease Diagnostic Laboratories (RADDLs).

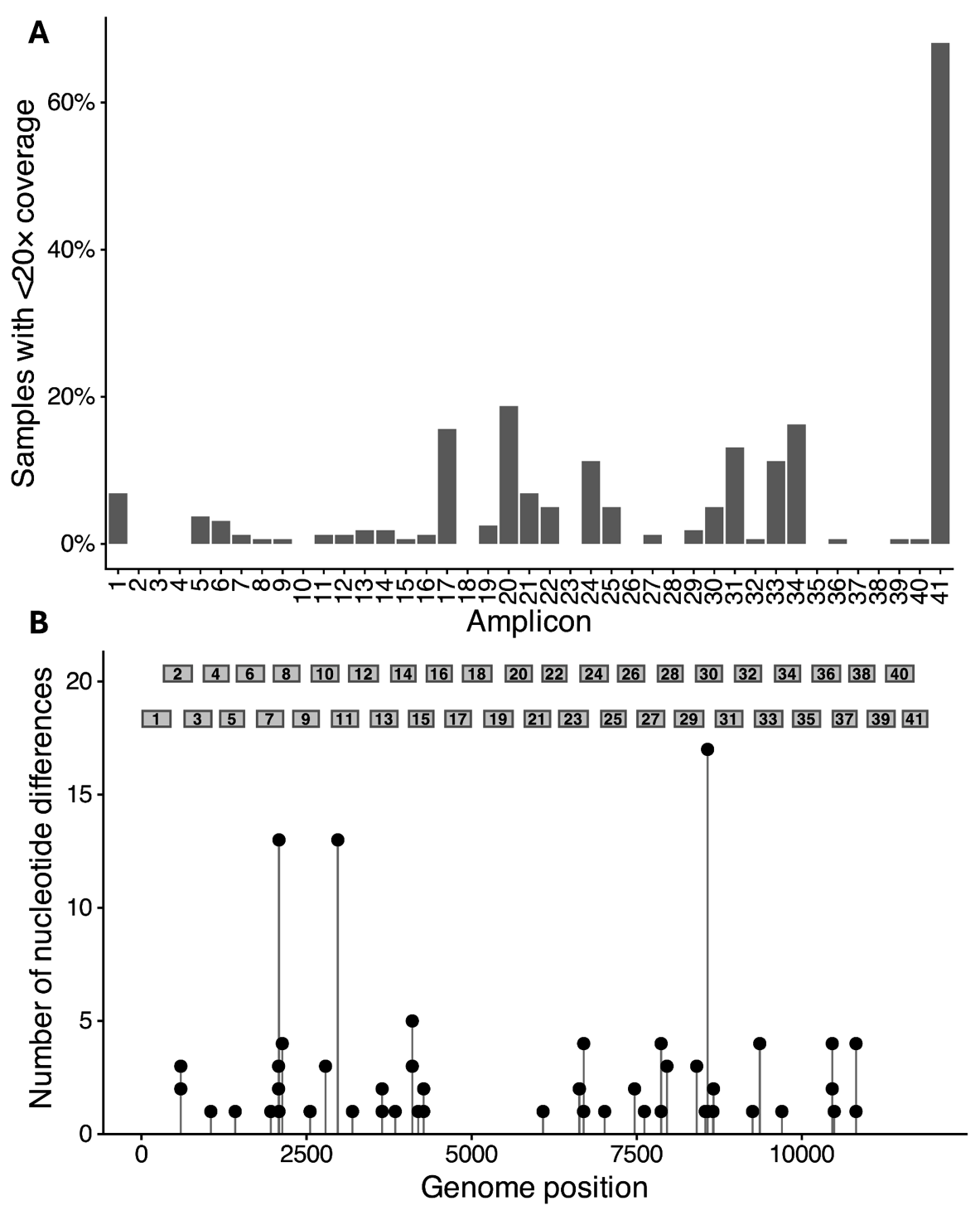

**Supplementary Figure 2. Amplicon failure and recurrent nucleotide differences across 160 rabies virus specimens. (**A) Bar plots show the proportion of specimens (%) that failed to achieve at least 20x coverage for each of the 41 amplicons spanning the rabies virus genome. (B) Lollipop plot shows the frequency and genomic position of unambiguous nucleotide differences between each sequenced specimen (FTA card or lateral flow device) and its matched baseline specimen (brain tissue). Amplicon regions are shown above the plot, with odd- and even-numbered amplicons staggered to indicate the two primer pools.

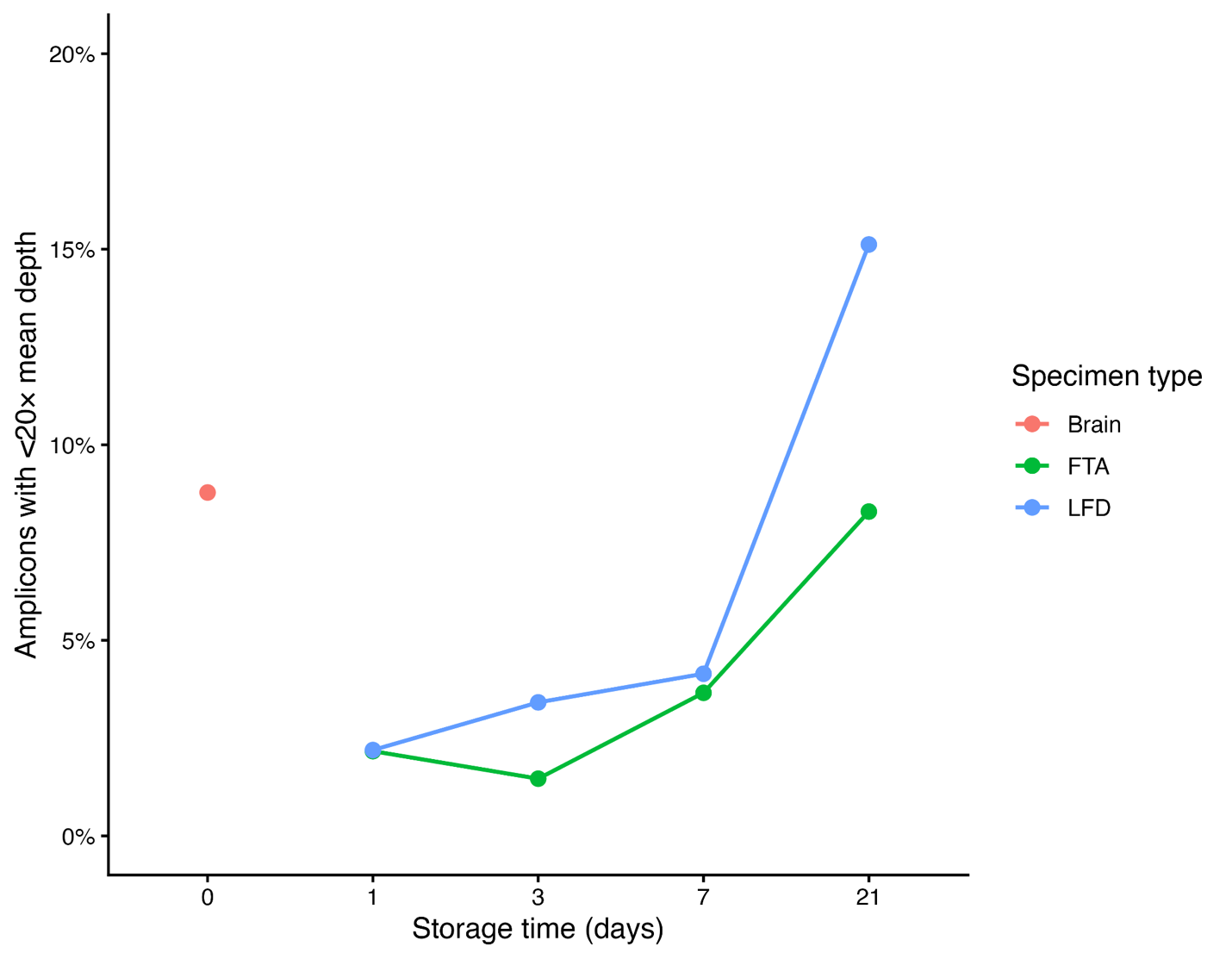

**Supplementary Figure 3. Amplicon failures over time for rabies brain, FTA-card and lateral flow device (LFD) specimens.** Plots are faceted by amplicon to show the proportion of specimens that failed to achieve 20x amplicon coverage. Brain tissue sequenced at day 0 was the baseline specimen, subsequent specimens represent FTA-cards and LFDs.

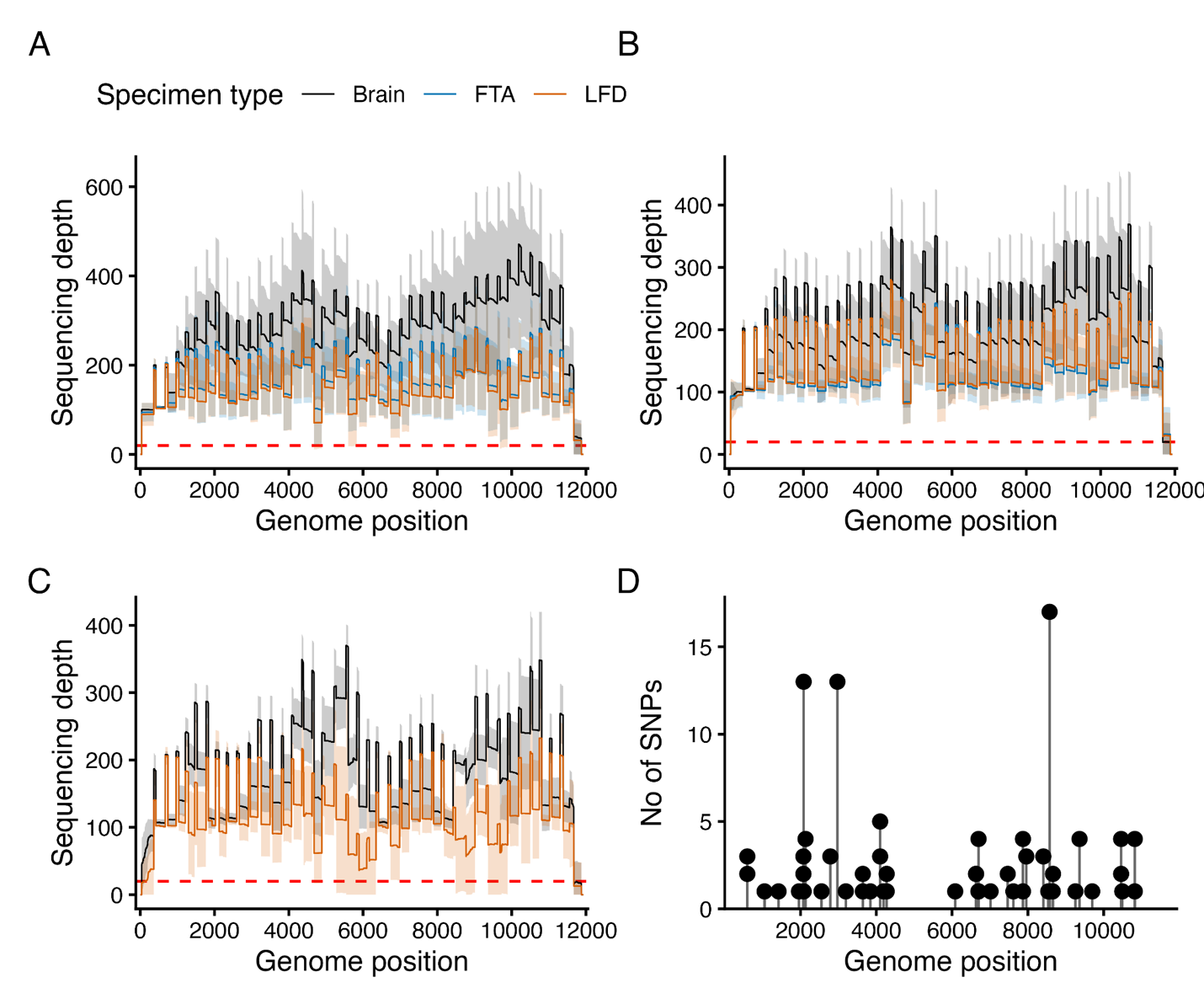

**Supplementary Figure 4. Sequencing read depth and nucleotide variation across the rabies virus genome.** Panels A-C show cohort-level summaries of sequencing read depth for (A) Cohort 1, (B) Cohort 2 and (C) Cohort 3 with the 20x read depth threshold indicated by a red dashed line. Panel D shows a lollipop plot of the overall frequency and genomic position of unambiguous nucleotide differences between each sequenced specimen (FTA card or lateral flow device) and its matched baseline specimen (brain tissue) across all cohorts.

**Supplementary Table 1.** List of rabies virus samples from domestic dogs in the Philippines used in this study.

| **Cohort** | **Study ID** | **Sample ID** | **Municipality** | **Province** | **Region** | **Date sample collection** | **Date sample received at RITM** |
| --- | --- | --- | --- | --- | --- | --- | --- |
| 1 | S1 | RADDL4B-25-046 | San Agustin | Romblon | MIMAROPA | 19-Jun-25 | 23-Sep-25 |
|  | S2 | RADDL4B-25-023 | San Agustin | Romblon | MIMAROPA | 15-May-25 | 23-Sep-25 |
|  | S3 | Z-25-053 | Silang | Cavite | Calabarzon | NA | 2-Oct-25 |
|  | S4 | Z-25-058 | Cavite City | Cavite | Calabarzon | NA | 8-Oct-25 |
|  | S5 | Z-25-061 | Cavite City | Cavite | Calabarzon | NA | 20-Oct-25 |
| 2 | N1 | Z-25-015 | Quezon City | Metro Manila | NCR | NA | 17-Feb-25 |
|  | N2 | Z-25-038 | Sta Rosa | Laguna | Calabarzon | NA | 9-Jul-25 |
|  | N3 | Z-26-007 | Biñan City | Laguna | Calabarzon | NA | 14-Jan-26 |
|  | N4 | Z-26-009 | Cavite City | Cavite | Calabarzon | NA | 21-Jan-26 |
|  | N5 | Z-26-012 | Las Piñas City | Metro Manila | NCR | NA | 30-Jan-26 |
|  | N6 | Z-26-017 | Cavite City | Cavite | Calabarzon | NA | 13-Feb-26 |
|  | N7 | Z-26-021 | Cavite City | Cavite | Calabarzon | NA | 23-Feb-26 |
|  | N8 | Z-26-023 | Cavite City | Cavite | Calabarzon | NA | 2-Mar-26 |
|  | N9 | Z-25-024 | Las Piñas City | Metro Manila | NCR | NA | 2-Mar-25 |
|  | N10 | Z-26-025 | Calamba City | Laguna | Calabarzon | NA | 6-Mar-26 |
|  | N11 | Z-26-031 | Muntinlupa City | Metro Manila | NCR | NA | 18-Mar-26 |
|  | N12 | Z-26-032 | Muntinlupa City | Metro Manila | NCR | NA | 18-Mar-26 |
|  | N13 | Z-26-035 | Dasmariñas City | Cavite | MIMAROPA | NA | 26-Mar-26 |
|  | N14 | RADDL4B-26-001 | San Jose | Occidental Mindoro | MIMAROPA | 24-Feb-26 | 24-Mar-26 |
|  | N15 | RADDL4B-26-005 | Magsaysay | Occidental Mindoro | MIMAROPA | 10-Mar-26 | 24-Mar-26 |
| 3 | - | RADDL4B-26-016 | Roxas | Oriental Mindoro | MIMAROPA | 21-May-26 | 6-Aug-26 |
|  | - | RADDL4B-26-017 | Pinamalayan | Oriental Mindoro | MIMAROPA | 17-Jun-26 | 6-Aug-26 |
|  | - | RADDL4B-26-019 | Rizal | Occidental Mindoro | MIMAROPA | 29-Jun-26 | 6-Aug-26 |
|  | - | RADDL4B-26-021 | Rizal | Occidental Mindoro | MIMAROPA | 8-Jul-26 | 6-Aug-26 |
|  | - | RADDL4B-26-022 | Pinamalayan | Oriental Mindoro | MIMAROPA | 4-Aug-26 | 6-Aug-26 |
|  | - | RADDL4B-26-024 | San Jose | Occidental Mindoro | MIMAROPA | 10-Jul-26 | 6-Aug-26 |
|  | - | RADDL4B-26-025 | Rizal | Occidental Mindoro | MIMAROPA | 16-Jul-26 | 6-Aug-26 |
|  | - | RADDL4B-26-027 | Rizal | Occidental Mindoro | MIMAROPA | 30-Jul-26 | 6-Aug-26 |

**Supplementary Table 2. Rabies virus specimen-level experimental conditions, real-time qPCR and sequencing results.** The table summarises the storage conditions and timepoints for all brain tissue, FTA-card and LFD specimens included in sequencing across the three study cohorts, together with corresponding real-time qPCR results and sequencing metrics.

| **Cohort** | **Specimen ID** | **Sample ID** | **Sample type** | **Temperature** | **Timepoint (days)** | **Quantification cycle (Cq)** | **Sequencing run ID** | **Mapped reads** | **Mean reads** | **Stdev reads** | **Bases Covered x20 (non-masked)** | **Genome coverage at ≥20× depth (%)** | **Lineage** |
| --- | --- | --- | --- | --- | --- | --- | --- | --- | --- | --- | --- | --- | --- |
| 1 | S1-D0 | S1 | Brain | NA | 0 | 23.27 | run1 | 18466.00 | 1364.13 | 386.08 | 11893.00 | 98.37 | SEA4_AI1 |
| 1 | S1-FTA-37-D1 | S1 | FTA | 37 | 1 | 18.68 | run2 | 47926.00 | 2256.68 | 1013.07 | 11886.00 | 98.61 | SEA4_AI1 |
| 1 | S1-FTA-37-D3 | S1 | FTA | 37 | 3 | 18.77 | run2 | 60685.00 | 3159.17 | 1254.93 | 11893.00 | 98.66 | SEA4_AI1 |
| 1 | S1-FTA-RT-D3 | S1 | FTA | RT | 3 | 20.3 | run2 | 37003.00 | 1956.22 | 757.98 | 11887.00 | 98.54 | SEA4_AI1 |
| 1 | S1-FTA-37-D7 | S1 | FTA | 37 | 7 | 19.01 | run2 | 80245.00 | 2754.61 | 1717.32 | 11886.00 | 97.12 | SEA4_AI1 |
| 1 | S1-FTA-RT-D7 | S1 | FTA | RT | 7 | 18.72 | run2 | 52286.00 | 2015.70 | 1008.94 | 11885.00 | 98.66 | SEA4_AI1 |
| 1 | S1-FTA-37-D21 | S1 | FTA | 37 | 21 | 21.02 | run2 | 50754.00 | 1833.05 | 1435.11 | 11361.00 | 78.42 | SEA4_AI1 |
| 1 | S1-FTA-RT-D21 | S1 | FTA | RT | 21 | 19.32 | run2 | 76121.00 | 2779.04 | 2152.07 | 11886.00 | 88.92 | SEA4_AI1 |
| 1 | S1-LFD-37-D1 | S1 | LFD | 37 | 1 | 23.14 | run2 | 47889.00 | 2751.34 | 1025.20 | 11893.00 | 98.62 | SEA4_AI1 |
| 1 | S1-LFD-RT-D1 | S1 | LFD | RT | 1 | 22.42 | run2 | 17097.00 | 1258.25 | 348.19 | 11893.00 | 98.36 | SEA4_AI1 |
| 1 | S1-LFD-37-D3 | S1 | LFD | 37 | 3 | 24.23 | run2 | 64939.00 | 2625.53 | 1536.64 | 11887.00 | 98.70 | SEA4_AI1 |
| 1 | S1-LFD-RT-D3 | S1 | LFD | RT | 3 | 23.37 | run2 | 68480.00 | 3703.54 | 1370.17 | 11894.00 | 98.57 | SEA4_AI1 |
| 1 | S1-LFD-37-D7 | S1 | LFD | 37 | 7 | 23.28 | run1 | 38773.00 | 1379.22 | 847.99 | 11885.00 | 98.44 | SEA4_AI1 |
| 1 | S1-LFD-RT-D7 | S1 | LFD | RT | 7 | 23.5 | run2 | 65968.00 | 3376.08 | 1551.52 | 11893.00 | 98.62 | SEA4_AI1 |
| 1 | S1-LFD-37-D21 | S1 | LFD | 37 | 21 | 24.46 | run2 | 49480.00 | 1762.18 | 1513.47 | 11318.00 | 88.88 | SEA4_AI1 |
| 1 | S1-LFD-RT-D21 | S1 | LFD | RT | 21 | 22.85 | run2 | 44234.00 | 1697.96 | 1338.16 | 11887.00 | 98.48 | SEA4_AI1 |
| 1 | S2-D0 | S2 | Brain | NA | 0 | 16.15 | run3 | 15510.00 | 1112.27 | 345.07 | 11915.00 | 97.90 | SEA4_R1.2.1 |
| 1 | S2-FTA-37-D1 | S2 | FTA | 37 | 1 | 14.25 | run3 | 21538.00 | 1165.80 | 508.92 | 11885.00 | 98.65 | SEA4_R1.2.1 |
| 1 | S2-FTA-RT-D1 | S2 | FTA | RT | 1 | 14.46 | run3 | 22550.00 | 1637.66 | 479.73 | 11886.00 | 98.59 | SEA4_R1.2.1 |
| 1 | S2-FTA-37-D3 | S2 | FTA | 37 | 3 | 14.21 | run3 | 34043.00 | 1406.40 | 792.03 | 11886.00 | 98.66 | SEA4_R1.2.1 |
| 1 | S2-FTA-RT-D3 | S2 | FTA | RT | 3 | 14.5 | run3 | 52804.00 | 2910.37 | 1150.87 | 11886.00 | 98.67 | SEA4_R1.2.1 |
| 1 | S2-FTA-37-D7 | S2 | FTA | 37 | 7 | 15.07 | run3 | 57795.00 | 2041.51 | 1497.65 | 11887.00 | 98.30 | SEA4_R1.2.1 |
| 1 | S2-FTA-RT-D7 | S2 | FTA | RT | 7 | 14.16 | run3 | 50548.00 | 2015.78 | 1161.46 | 11887.00 | 98.65 | SEA4_R1.2.1 |
| 1 | S2-FTA-37-D21 | S2 | FTA | 37 | 21 | 15.62 | run3 | 43043.00 | 1561.08 | 1292.56 | 11885.00 | 90.97 | SEA4_R1.2.1 |
| 1 | S2-FTA-RT-D21 | S2 | FTA | RT | 21 | 14.88 | run4 | 44979.00 | 1664.04 | 1274.52 | 11884.00 | 95.07 | SEA4_R1.2.1 |
| 1 | S2-LFD-37-D1 | S2 | LFD | 37 | 1 | 22.26 | run3 | 39526.00 | 1575.32 | 1074.78 | 11887.00 | 98.67 | SEA4_R1.2.1 |
| 1 | S2-LFD-RT-D1 | S2 | LFD | RT | 1 | 21.03 | run3 | 33012.00 | 1464.90 | 759.51 | 11915.00 | 98.72 | SEA4_R1.2.1 |
| 1 | S2-LFD-37-D3 | S2 | LFD | 37 | 3 | 21.96 | run3 | 55146.00 | 2102.18 | 1493.58 | 11887.00 | 98.68 | SEA4_R1.2.1 |
| 1 | S2-LFD-RT-D3 | S2 | LFD | RT | 3 | 21.17 | run3 | 24375.00 | 1070.38 | 599.64 | 11892.00 | 98.68 | SEA4_R1.2.1 |
| 1 | S2-LFD-37-D7 | S2 | LFD | 37 | 7 | 21.45 | run3 | 35122.00 | 1300.18 | 1097.40 | 11885.00 | 95.75 | SEA4_R1.2.1 |
| 1 | S2-LFD-RT-D7 | S2 | LFD | RT | 7 | 20.83 | run3 | 32238.00 | 1303.38 | 785.49 | 11887.00 | 98.58 | SEA4_R1.2.1 |
| 1 | S2-LFD-37-D21 | S2 | LFD | 37 | 21 | 22.57 | run3 | 24075.00 | 912.99 | 820.00 | 11154.00 | 85.58 | SEA4_R1.2.1 |
| 1 | S2-LFD-RT-D21 | S2 | LFD | RT | 21 | 21.5 | run3 | 23916.00 | 872.41 | 782.05 | 11688.00 | 95.04 | SEA4_R1.2.1 |
| 1 | S3-D0 | S3 | Brain | NA | 0 | 17.6 | run3 | 6469.00 | 478.19 | 157.61 | 11675.00 | 96.72 | SEA4_Y1.1.1 |
| 1 | S3-FTA-37-D1 | S3 | FTA | 37 | 1 | 17.51 | run3 | 37984.00 | 1776.91 | 895.05 | 11885.00 | 98.67 | SEA4_Y1.1.1 |
| 1 | S3-FTA-RT-D1 | S3 | FTA | RT | 1 | 17.75 | run3 | 17549.00 | 946.21 | 433.69 | 11692.00 | 97.09 | SEA4_Y1.1.1 |
| 1 | S3-FTA-37-D3 | S3 | FTA | 37 | 3 | 18.07 | run3 | 30337.00 | 1124.99 | 648.07 | 11678.00 | 95.53 | SEA4_Y1.1.1 |
| 1 | S3-FTA-RT-D3 | S3 | FTA | RT | 3 | 17.92 | run3 | 20925.00 | 1038.98 | 459.92 | 11677.00 | 97.00 | SEA4_Y1.1.1 |
| 1 | S3-FTA-37-D7 | S3 | FTA | 37 | 7 | 16.58 | run3 | 48248.00 | 1698.32 | 1067.14 | 11687.00 | 96.80 | SEA4_Y1.1.1 |
| 1 | S3-FTA-RT-D7 | S3 | FTA | RT | 7 | 16.18 | run3 | 27637.00 | 1111.49 | 621.00 | 11680.00 | 97.00 | SEA4_Y1.1.1 |
| 1 | S3-FTA-37-D21 | S3 | FTA | 37 | 21 | 17.17 | run3 | 32430.00 | 1148.65 | 868.44 | 11788.00 | 93.50 | SEA4_Y1.1.1 |
| 1 | S3-FTA-RT-D21 | S3 | FTA | RT | 21 | 17.43 | run3 | 34229.00 | 1357.62 | 1120.13 | 11693.00 | 88.55 | SEA4_Y1.1.1 |
| 1 | S3-LFD-37-D1 | S3 | LFD | 37 | 1 | 21.98 | run3 | 21200.00 | 956.53 | 511.60 | 11678.00 | 97.02 | SEA4_Y1.1.1 |
| 1 | S3-LFD-RT-D1 | S3 | LFD | RT | 1 | 21.85 | run3 | 32041.00 | 1399.26 | 803.90 | 11692.00 | 97.01 | SEA4_Y1.1.1 |
| 1 | S3-LFD-37-D3 | S3 | LFD | 37 | 3 | 22.25 | run3 | 44019.00 | 1627.16 | 1143.48 | 11692.00 | 96.85 | SEA4_Y1.1.1 |
| 1 | S3-LFD-RT-D3 | S3 | LFD | RT | 3 | 22.1 | run3 | 24663.00 | 1049.02 | 596.49 | 11685.00 | 96.92 | SEA4_Y1.1.1 |
| 1 | S3-LFD-37-D7 | S3 | LFD | 37 | 7 | 22.19 | run3 | 29876.00 | 1094.43 | 789.38 | 11682.00 | 93.62 | SEA4_Y1.1.1 |
| 1 | S3-LFD-RT-D7 | S3 | LFD | RT | 7 | 22.57 | run3 | 28523.00 | 1130.80 | 722.72 | 11692.00 | 95.66 | SEA4_Y1.1.1 |
| 1 | S3-LFD-37-D21 | S3 | LFD | 37 | 21 | 24.32 | run3 | 10115.00 | 379.22 | 356.28 | 10330.00 | 81.31 | SEA4_Y1.1.1 |
| 1 | S3-LFD-RT-D21 | S3 | LFD | RT | 21 | 22.87 | run3 | 35013.00 | 1265.65 | 976.97 | 11515.00 | 86.19 | SEA4_Y1.1.1 |
| 1 | S4-D0 | S4 | Brain | NA | 0 | 15.16 | run1 | 1845.00 | 124.23 | 46.52 | 11670.00 | 96.89 | SEA4_AL1 |
| 1 | S4-FTA-37-D1 | S4 | FTA | 37 | 1 | 17.63 | run1 | 9881.00 | 443.22 | 225.54 | 11881.00 | 96.97 | SEA4_AL1 |
| 1 | S4-FTA-RT-D1 | S4 | FTA | RT | 1 | 18.39 | run1 | 6414.00 | 306.64 | 153.39 | 11882.00 | 96.81 | SEA4_AL1 |
| 1 | S4-FTA-37-D3 | S4 | FTA | 37 | 3 | 18.76 | run1 | 16001.00 | 604.93 | 401.92 | 11691.00 | 94.99 | SEA4_AL1 |
| 1 | S4-FTA-RT-D3 | S4 | FTA | RT | 3 | 17.91 | run1 | 10677.00 | 495.93 | 251.59 | 11883.00 | 96.86 | SEA4_AL1 |
| 1 | S4-FTA-37-D7 | S4 | FTA | 37 | 7 | 21.01 | run1 | 8655.00 | 320.97 | 272.53 | 10951.00 | 78.05 | SEA4_AL1 |
| 1 | S4-FTA-RT-D7 | S4 | FTA | RT | 7 | 19.09 | run1 | 7980.00 | 293.58 | 232.06 | 11669.00 | 90.46 | SEA4_AL1 |
| 1 | S4-FTA-37-D21 | S4 | FTA | 37 | 21 | 24.39 | run1 | 3029.00 | 110.84 | 110.30 | 9102.00 | 70.45 | SEA4_AL1 |
| 1 | S4-FTA-RT-D21 | S4 | FTA | RT | 21 | 21.19 | run1 | 6827.00 | 243.20 | 228.79 | 10109.00 | 80.35 | SEA4_AL1 |
| 1 | S4-LFD-37-D1 | S4 | LFD | 37 | 1 | 21.33 | run1 | 10120.00 | 441.97 | 248.35 | 11696.00 | 96.73 | SEA4_AL1 |
| 1 | S4-LFD-RT-D1 | S4 | LFD | RT | 1 | 20.8 | run1 | 7079.00 | 354.58 | 165.05 | 11888.00 | 96.85 | SEA4_AL1 |
| 1 | S4-LFD-37-D3 | S4 | LFD | 37 | 3 | 22.09 | run1 | 8928.00 | 324.03 | 275.91 | 11312.00 | 86.64 | SEA4_AL1 |
| 1 | S4-LFD-RT-D3 | S4 | LFD | RT | 3 | 21.56 | run1 | 8559.00 | 355.32 | 210.45 | 11678.00 | 96.93 | SEA4_AL1 |
| 1 | S4-LFD-37-D7 | S4 | LFD | 37 | 7 | 22.66 | run1 | 9776.00 | 370.64 | 307.81 | 11292.00 | 83.78 | SEA4_AL1 |
| 1 | S4-LFD-RT-D7 | S4 | LFD | RT | 7 | NA | run1 | 9341.00 | 368.80 | 310.22 | 11492.00 | 86.07 | SEA4_AL1 |
| 1 | S4-LFD-37-D21 | S4 | LFD | 37 | 21 | 23.75 | run1 | 1992.00 | 75.27 | 74.30 | 8685.00 | 64.22 | SEA4_AL1 |
| 1 | S4-LFD-RT-D21 | S4 | LFD | RT | 21 | 24.15 | run1 | 2999.00 | 111.04 | 106.97 | 9515.00 | 65.02 | SEA4_AL1 |
| 1 | S5-D0 | S5 | Brain | NA | 0 | 18.28 | run4 | 23790.00 | 2065.30 | 630.72 | 11893.00 | 98.15 | SEA4_AD1.2.1 |
| 1 | S5-FTA-37-D1 | S5 | FTA | 37 | 1 | 17.3 | run4 | 36382.00 | 2353.02 | 923.35 | 11892.00 | 98.66 | SEA4_AD1.2.1 |
| 1 | S5-FTA-RT-D1 | S5 | FTA | RT | 1 | 17.01 | run4 | 23065.00 | 2085.66 | 564.86 | 11892.00 | 96.72 | SEA4_AD1.2.1 |
| 1 | S5-FTA-37-D3 | S5 | FTA | 37 | 3 | 17.71 | run4 | 30135.00 | 1345.60 | 740.21 | 11825.00 | 96.96 | SEA4_AD1.2.1 |
| 1 | S5-FTA-RT-D3 | S5 | FTA | RT | 3 | 17.21 | run4 | 19083.00 | 1204.60 | 577.42 | 11886.00 | 96.89 | SEA4_AD1.2.1 |
| 1 | S5-FTA-37-D7 | S5 | FTA | 37 | 7 | 19.79 | run4 | 28446.00 | 1154.74 | 775.36 | 11842.00 | 95.22 | SEA4_AD1.2.1 |
| 1 | S5-FTA-RT-D7 | S5 | FTA | RT | 7 | 18.06 | run4 | 24221.00 | 1264.88 | 646.73 | 11837.00 | 96.89 | SEA4_AD1.2.1 |
| 1 | S5-FTA-37-D21 | S5 | FTA | 37 | 21 | 21.08 | run4 | 14834.00 | 589.95 | 455.07 | 11676.00 | 81.05 | SEA4_AD1.2.1 |
| 1 | S5-FTA-RT-D21 | S5 | FTA | RT | 21 | 20.04 | run4 | 27941.00 | 1082.97 | 791.74 | 11693.00 | 95.05 | SEA4_AD1.2.1 |
| 1 | S5-LFD-37-D1 | S5 | LFD | 37 | 1 | 24.41 | run4 | 33698.00 | 1636.48 | 947.57 | 11892.00 | 98.57 | SEA4_AD1.2.1 |
| 1 | S5-LFD-RT-D1 | S5 | LFD | RT | 1 | 23.5 | run4 | 33645.00 | 1704.85 | 896.35 | 11892.00 | 96.87 | SEA4_AD1.2.1 |
| 1 | S5-LFD-37-D3 | S5 | LFD | 37 | 3 | 25.19 | run4 | 22326.00 | 1049.50 | 611.64 | 11884.00 | 96.91 | SEA4_AD1.2.1 |
| 1 | S5-LFD-RT-D3 | S5 | LFD | RT | 3 | 23.51 | run4 | 21901.00 | 1185.54 | 639.86 | 11892.00 | 98.59 | SEA4_AD1.2.1 |
| 1 | S5-LFD-37-D7 | S5 | LFD | 37 | 7 | 24.8 | run4 | 21118.00 | 920.49 | 677.37 | 11758.00 | 96.88 | SEA4_AD1.2.1 |
| 1 | S5-LFD-RT-D7 | S5 | LFD | RT | 7 | 23.36 | run4 | 18274.00 | 1009.48 | 532.63 | 11892.00 | 98.63 | SEA4_AD1.2.1 |
| 1 | S5-LFD-37-D21 | S5 | LFD | 37 | 21 | 26.03 | run4 | 11893.00 | 476.40 | 384.49 | 11132.00 | 84.17 | SEA4_AD1.2.1 |
| 1 | S5-LFD-RT-D21 | S5 | LFD | RT | 21 | 25.7 | run4 | 14989.00 | 600.19 | 484.43 | 11678.00 | 89.17 | SEA4_AD1.2.1 |
| 2 | N1-D0 | N1 | Brain | NA | 0 | 17.08 | run9 | 108549.00 | 5004.29 | 2542.19 | 11693.00 | 96.87 | SEA4_AC1.1.1 |
| 2 | N1-FTA-D3 | N1 | FTA | RT | 3 | 18.66 | run9 | 181554.00 | 6657.00 | 4376.58 | 11700.00 | 94.33 | SEA4_AC1.1.1 |
| 2 | N1-FTA-D7 | N1 | FTA | RT | 7 | 17.32 | run9 | 90350.00 | 3374.31 | 2443.88 | 11693.00 | 91.32 | SEA4_AC1.1.1 |
| 2 | N1-LFD-D3 | N1 | LFD | RT | 3 | 21.94 | run9 | 79333.00 | 2826.11 | 2472.13 | 11692.00 | 92.82 | SEA4_AC1.1.1 |
| 2 | N1-LFD-D7 | N1 | LFD | RT | 7 | 22.88 | run9 | 86459.00 | 3127.85 | 2682.83 | 11684.00 | 93.44 | SEA4_AC1.1.1 |
| 2 | N-12-D0 | N12 | Brain | NA | 0 | NA | run5 | 20650.00 | 938.21 | 500.70 | 11885.00 | 96.72 | SEA4_P1.2.2 |
| 2 | N-12-FTA-D3 | N12 | FTA | RT | 3 | 21.45 | run5 | 33521.00 | 1385.75 | 769.33 | 11886.00 | 98.56 | SEA4_P1.2.2 |
| 2 | N-12-FTA-D7 | N12 | FTA | RT | 7 | 21.58 | run5 | 19973.00 | 685.47 | 513.44 | 11676.00 | 96.55 | SEA4_P1.2.2 |
| 2 | N-12-LFD-D3 | N12 | LFD | RT | 3 | 27.17 | run5 | 21768.00 | 911.63 | 538.91 | 11893.00 | 98.20 | SEA4_P1.2.2 |
| 2 | N-12-LFD-D7 | N12 | LFD | RT | 7 | 28.24 | run5 | 40471.00 | 1468.78 | 1014.23 | 11886.00 | 94.67 | SEA4_P1.2.2 |
| 2 | N-13-D0 | N13 | Brain | NA | 0 | 23.52 | run5 | 12185.00 | 873.23 | 269.13 | 11892.00 | 98.30 | SEA4_AK1 |
| 2 | N-13-FTA-D3 | N13 | FTA | RT | 3 | 24.05 | run5 | 25319.00 | 1057.32 | 557.51 | 11886.00 | 98.41 | SEA4_AK1 |
| 2 | N-13-FTA-D7 | N13 | FTA | RT | 7 | 23.94 | run5 | 40410.00 | 1414.18 | 999.61 | 11886.00 | 98.39 | SEA4_AK1 |
| 2 | N-13-LFD-D3 | N13 | LFD | RT | 3 | 28.39 | run5 | 25746.00 | 1148.55 | 602.52 | 11893.00 | 98.47 | SEA4_AK1 |
| 2 | N-13-LFD-D7 | N13 | LFD | RT | 7 | 27.4 | run5 | 30403.00 | 1120.92 | 748.47 | 11885.00 | 98.43 | SEA4_AK1 |
| 2 | N-14-D0 | N14 | Brain | NA | 0 | NA | run11 | 143209.00 | 5540.30 | 6310.73 | 11671.00 | 95.78 | SEA4_AJ1 |
| 2 | N-14-FTA-D3 | N14 | FTA | RT | 3 | 21.45 | run11 | 122060.00 | 5048.62 | 5451.71 | 11692.00 | 95.33 | SEA4_AJ1 |
| 2 | N-14-FTA-D7 | N14 | FTA | RT | 7 | 26.35 | run11 | 139699.00 | 5499.51 | 5498.12 | 11342.00 | 75.10 | SEA4_AJ1 |
| 2 | N-14-LFD-D3 | N14 | LFD | RT | 3 | 21.63 | run11 | 139991.00 | 5773.86 | 6853.16 | 11809.00 | 95.50 | SEA4_AJ1 |
| 2 | N-14-LFD-D7 | N14 | LFD | RT | 7 | 26.86 | run11 | 136004.00 | 5338.82 | 6655.07 | 11579.00 | 90.15 | SEA4_AJ1 |
| 2 | N-15-D0 | N15 | Brain | NA | 0 | NA | run9 | 108125.00 | 4288.64 | 3207.21 | 11886.00 | 98.48 | SEA4_P1.1.1 |
| 2 | N-15-FTA-D3 | N15 | FTA | RT | 3 | 19.33 | run9 | 79409.00 | 3463.68 | 1837.49 | 11887.00 | 98.41 | SEA4_P1.1.1 |
| 2 | N-15-FTA-D7 | N15 | FTA | RT | 7 | 26.13 | run9 | 42764.00 | 1549.74 | 1153.74 | 11886.00 | 98.17 | SEA4_P1.1.1 |
| 2 | N-15-LFD-D3 | N15 | LFD | RT | 3 | 19.63 | run9 | 73802.00 | 2910.44 | 1961.70 | 11888.00 | 98.28 | SEA4_P1.1.1 |
| 2 | N-15-LFD-D7 | N15 | LFD | RT | 7 | 25.72 | run9 | 103916.00 | 3834.82 | 3342.12 | 11886.00 | 93.19 | SEA4_P1.1.1 |
| 2 | N2-D0 | N2 | Brain | NA | 0 | 19.1 | run9 | 71879.00 | 3290.82 | 1876.66 | 11682.00 | 96.80 | SEA4_AF1 |
| 2 | N2-FTA-D3 | N2 | FTA | RT | 3 | 17.28 | run9 | 105147.00 | 4209.29 | 2994.75 | 11693.00 | 96.81 | SEA4_AF1 |
| 2 | N2-FTA-D7 | N2 | FTA | RT | 7 | 16.87 | run9 | 172682.00 | 6334.26 | 4294.39 | 11693.00 | 93.94 | SEA4_AF1 |
| 2 | N2-LFD-D3 | N2 | LFD | RT | 3 | 20.41 | run9 | 142467.00 | 5563.73 | 3823.57 | 11886.00 | 98.48 | SEA4_AF1 |
| 2 | N2-LFD-D7 | N2 | LFD | RT | 7 | 21.8 | run9 | 133655.00 | 4826.54 | 4454.84 | 11884.00 | 90.94 | SEA4_AF1 |
| 2 | N3-D0 | N3 | Brain | NA | 0 | 18.38 | run9 | 55968.00 | 3169.17 | 1421.49 | 11683.00 | 96.65 | SEA4_AF1.1 |
| 2 | N3-FTA-D3 | N3 | FTA | RT | 3 | 16.08 | run9 | 107233.00 | 4414.05 | 2786.85 | 11689.00 | 96.85 | SEA4_AF1.1 |
| 2 | N3-FTA-D7 | N3 | FTA | RT | 7 | 15.05 | run9 | 94279.00 | 3510.59 | 2880.81 | 11693.00 | 93.79 | SEA4_AF1.1 |
| 2 | N3-LFD-D3 | N3 | LFD | RT | 3 | 18.78 | run9 | 107288.00 | 4369.57 | 2602.82 | 11685.00 | 96.68 | SEA4_AF1.1 |
| 2 | N3-LFD-D7 | N3 | LFD | RT | 7 | 19.8 | run9 | 105678.00 | 3911.25 | 3321.62 | 11693.00 | 93.77 | SEA4_AF1.1 |
| 2 | N4-D0 | N4 | Brain | NA | 0 | 18.35 | run9 | 186810.00 | 7058.72 | 5427.69 | 11712.00 | 93.93 | SEA4_AD1.2.1 |
| 2 | N4-FTA-D3 | N4 | FTA | RT | 3 | 20.04 | run9 | 111251.00 | 4064.73 | 2632.62 | 11693.00 | 95.37 | SEA4_AD1.2.1 |
| 2 | N4-FTA-D7 | N4 | FTA | RT | 7 | 19.85 | run9 | 110546.00 | 4019.62 | 3199.33 | 11693.00 | 89.97 | SEA4_AD1.2.1 |
| 2 | N4-LFD-D3 | N4 | LFD | RT | 3 | 21.04 | run9 | 70196.00 | 2783.14 | 1918.23 | 11693.00 | 96.77 | SEA4_AD1.2.1 |
| 2 | N4-LFD-D7 | N4 | LFD | RT | 7 | 22.82 | run9 | 79797.00 | 2879.23 | 2402.55 | 11693.00 | 96.65 | SEA4_AD1.2.1 |
| 2 | N-5-D0 | N5 | Brain | NA | 0 | 20.17 | run9 | 47256.00 | 2218.91 | 1252.35 | 11710.00 | 96.79 | SEA4_AD1.2 |
| 2 | N-5-FTA-D3 | N5 | FTA | RT | 3 | 22.65 | run9 | 124241.00 | 4536.97 | 3254.91 | 11709.00 | 93.64 | SEA4_AD1.2 |
| 2 | N-5-FTA-D7 | N5 | FTA | RT | 7 | 21.95 | run9 | 154132.00 | 5512.16 | 3913.87 | 11887.00 | 98.33 | SEA4_AD1.2 |
| 2 | N-5-LFD-D3 | N5 | LFD | RT | 3 | 22.98 | run9 | 123655.00 | 4424.57 | 3766.22 | 11692.00 | 92.57 | SEA4_AD1.2 |
| 2 | N-5-LFD-D7 | N5 | LFD | RT | 7 | 24.93 | run9 | 133050.00 | 4785.79 | 4136.28 | 11702.00 | 92.26 | SEA4_AD1.2 |
| 2 | N-7-D0 | N7 | Brain | NA | 0 | 23.46 | run7 | 65257.00 | 2955.69 | 1623.73 | 11886.00 | 98.75 | SEA4_AL1 |
| 2 | N-7-FTA-D3 | N7 | FTA | RT | 3 | 24.57 | run7 | 49292.00 | 1815.30 | 1926.09 | 11693.00 | 97.02 | SEA4_AL1 |
| 2 | N-7-FTA-D7 | N7 | FTA | RT | 7 | 25.31 | run7 | 20883.00 | 725.77 | 1002.77 | 11499.00 | 91.03 | SEA4_AL1 |
| 2 | N-7-LFD-D3 | N7 | LFD | RT | 3 | 29.56 | run7 | 31547.00 | 1104.43 | 1472.48 | 11512.00 | 89.40 | SEA4_AL1 |
| 2 | N-7-LFD-D7 | N7 | LFD | RT | 7 | 31.33 | run7 | 15854.00 | 581.55 | 740.17 | 10380.00 | 80.17 | SEA4_AL1 |
| 2 | N-8-D0 | N8 | Brain | NA | 0 | 20.7 | run7 | 36709.00 | 2376.77 | 994.45 | 11892.00 | 96.87 | SEA4_AL1 |
| 2 | N-8-FTA-D3 | N8 | FTA | RT | 3 | 20.15 | run7 | 20665.00 | 840.83 | 598.73 | 11675.00 | 97.10 | SEA4_AL1 |
| 2 | N-8-FTA-D7 | N8 | FTA | RT | 7 | 20.01 | run7 | 27360.00 | 973.96 | 1179.77 | 11677.00 | 92.87 | SEA4_AL1 |
| 2 | N-8-LFD-D3 | N8 | LFD | RT | 3 | 24.58 | run7 | 38260.00 | 1617.53 | 1174.07 | 11887.00 | 98.79 | SEA4_AL1 |
| 2 | N-8-LFD-D7 | N8 | LFD | RT | 7 | 24.03 | run7 | 42624.00 | 1611.01 | 1467.43 | 11885.00 | 98.63 | SEA4_AL1 |
| 2 | N-9-D0 | N9 | Brain | NA | 0 | 19.68 | run7 | 33250.00 | 2122.41 | 1050.25 | 11887.00 | 98.56 | SEA4_AK1 |
| 2 | N-9-FTA-D3 | N9 | FTA | RT | 3 | 21.6 | run7 | 26367.00 | 1033.95 | 863.62 | 11887.00 | 98.90 | SEA4_AK1 |
| 2 | N-9-FTA-D7 | N9 | FTA | RT | 7 | 20.98 | run7 | 41570.00 | 1552.31 | 1759.94 | 11887.00 | 97.23 | SEA4_AK1 |
| 2 | N-9-LFD-D3 | N9 | LFD | RT | 3 | 22.34 | run7 | 73011.00 | 3888.06 | 1588.13 | 11894.00 | 98.73 | SEA4_AK1 |
| 2 | N-9-LFD-D7 | N9 | LFD | RT | 7 | 21.35 | run7 | 31243.00 | 1144.99 | 1155.79 | 11884.00 | 95.70 | SEA4_AK1 |
| 3 | NA | RADDL4B-2026-016 | Brain | Field | NA | NA | run11 | 66618.00 | 3081.17 | 2342.12 | 11689.00 | 97.10 | SEA4_P1.1.1 |
| 3 | NA | RADDL4B-2026-016-LFD | LFD | Field | 77 | NA | run11 | 5372.00 | 185.20 | 258.27 | 7740.00 | 52.83 | SEA4_P1.1.1 |
| 3 | NA | RADDL4B-2026-017 | Brain | Field | NA | NA | run11 | 104445.00 | 4362.34 | 5108.58 | 11755.00 | 97.10 | SEA4_P1.1.1 |
| 3 | NA | RADDL4B-2026-017-LFD | LFD | Field | 50 | NA | run11 | 22692.00 | 848.34 | 1255.69 | 9042.00 | 71.57 | SEA4_P1.1.1 |
| 3 | NA | RADDL4B-2026-019 | Brain | Field | NA | NA | run11 | 69503.00 | 3211.63 | 2577.38 | 11637.00 | 95.31 | SEA4_AJ1 |
| 3 | NA | RADDL4B-2026-019-LFD | LFD | Field | 35 | NA | run11 | 53613.00 | 1942.37 | 2474.19 | 10655.00 | 80.22 | SEA4_AJ1 |
| 3 | NA | RADDL4B-2026-021 | Brain | Field | NA | NA | run11 | 44268.00 | 2143.16 | 1460.56 | 11823.00 | 98.10 | SEA4_AJ1 |
| 3 | NA | RADDL4B-2026-021-LFD | LFD | Field | 29 | NA | run11 | 44401.00 | 1633.36 | 2330.61 | 11027.00 | 81.05 | SEA4_AJ1 |
| 3 | NA | RADDL4B-2026-022 | Brain | Field | NA | NA | run11 | 71526.00 | 3921.46 | 2703.04 | 11803.00 | 97.04 | SEA4_P1.1.1 |
| 3 | NA | RADDL4B-2026-022-LFD | LFD | Field | 1 | NA | run11 | 73511.00 | 2677.11 | 4593.29 | 10764.00 | 79.57 | SEA4_P1.1.1 |
| 3 | NA | RADDL4B-2026-024 | Brain | Field | NA | NA | run11 | 135104.00 | 5909.98 | 5071.56 | 11802.00 | 97.11 | SEA4_AJ1 |
| 3 | NA | RADDL4B-2026-024-LFD | LFD | Field | 23 | NA | run11 | 69129.00 | 2620.31 | 3681.07 | 11513.00 | 83.00 | SEA4_AJ1 |
| 3 | NA | RADDL4B-2026-025 | Brain | Field | NA | NA | run11 | 45250.00 | 2031.67 | 1865.62 | 11577.00 | 75.00 | SEA4_AJ1 |
| 3 | NA | RADDL4B-2026-025-LFD | LFD | Field | 20 | NA | run11 | 73321.00 | 3105.01 | 3004.27 | 11685.00 | 97.13 | SEA4_AJ1 |
| 3 | NA | RADDL4B-2026-027 | Brain | Field | NA | NA | run11 | 67897.00 | 3780.54 | 2448.53 | 11681.00 | 97.02 | SEA4_AJ1 |
| 3 | NA | RADDL4B-2026-027-LFD | LFD | Field | 1 | NA | run11 | 96225.00 | 4525.11 | 3213.26 | 11887.00 | 98.78 | SEA4_AJ1 |
